# *Ustilago maydis* disrupts carbohydrate signaling networks to induce hypertrophy in host cells

**DOI:** 10.1101/2024.11.22.624849

**Authors:** Yoon Joo Lee, Sara Christina Stolze, Georgios Saridis, Malaika K. Ebert, Hirofumi Nakagami, Gunther Doehlemann

**Author notes:** Department of Plant Pathology, North Dakota State University, NDSU Department 7660, P.O. Box 6050, Fargo, ND, 58108-6050 USA.

## Abstract

*Ustilago maydis* is a biotrophic fungus infecting maize, secreting effector proteins to manipulate host cellular processes and create nutrient-rich environments for its growth. Three effectors, Hap1-3 (hypertrophy-associated proteins), were identified as virulence factors promoting hypertrophic mesophyll tumor cells (HTT). Immunoprecipitation and mass spectrometry revealed interactions among Hap effectors, suggesting potential effector complex formation. CRISPR-Cas9 triple knockout of hap1-3 demonstrated Hap1’s role in HTT formation. Infection assays identified Hap1 as a key virulence factor interacting with maize Snf1-related kinase 1 (SnRK1), a central energy regulator. RNA-seq analysis showed that Hap1 promotes cell cycle and starch biosynthesis genes, while CR-*hap1* induced defense-related WRKY transcription factors. Phosphoproteomics revealed increased phosphorylation of SnRK1 and metabolic enzymes during SG200 infection. Our findings support a model where *U. maydis* induces hypertrophy through Hap1, targeting the SnRK1α subunit to prevent SnRK1 inhibition by high trehalose-6-phosphate (T6P), disrupting the antagonistic relationship between T6P and SnRK1. This reprograms host transcription to enhance starch metabolism and induce endoreduplication, leading to hypertrophy induction, while suppressing sugar-induced immune signaling.

## Introduction

Smut fungi are one of the largest groups of biotrophic plant pathogens, infecting a wide range of cereal plants ^1^. Most smuts spread disease systemically through the vascular system, causing symptoms exclusive to the inflorescences ^2,3^. However, *Ustilago maydis* forms local tumors in all aerial parts of maize ^4^. To infect maize organs, which vary substantially in their development and physiology, *U. maydis* secretes a cocktail of effectors in a spatiotemporal and organ-specific manner, promoting host colonization and suppressing plant immune responses ^5,6^. *In vivo* visualization of *U. maydis-*induced tumors showed that newly divided bundle sheath cells transform into hyperplasic tumor (HPT) cells, while mesophyll cells enlarge and become hypertrophic tumor (HTT) cells due to endoreduplication, a cellular mechanism characterized by DNA replication without cell division, resulting in increased nuclear DNA content ^7,8^. Targeted transcriptomic analysis using laser-captured microdissection identified specific effector gene expressions associated with HPT and HTT formation, highlighting the cell-type specific role of effectors in tumor formation ^7^. For example, See1 (Seedling efficient effector 1) interacts with ZmSGT1, a cell cycle transition regulator, to reactivate DNA synthesis essential for cell division. Moreover, Sts2 (Small tumor on seedlings 2) interacts with ZmNECAP1, a plant transcriptional activator, activating leaf development regulators ^8^. However, effectors involved in HTT formation, and their underlying molecular mechanisms remain uncharacterized.

*U. maydis* relies on living tissues for survival, invading the intercellular spaces by breaking down the loosened cell wall to extract nutrients from the host ^9^. Upon infection, it reprograms plant metabolic processes, redirecting starch and stimulating the accumulation of hexoses, sucrose, and sugar phosphates such as trehalose-6-phosphate (T6P) and glucose-6-phosphate (G6P) toward developing tumors, thereby transforming them into strong nutrient sinks ^10–13^. Notably, starch granules begin to excessively accumulate as early as 2 dpi, particularly in mesophyll HTT cells, which are atypical sites for starch storage in C4 plants ^7,14^. Moreover, several transcriptomic analyses have also shown that the infection leads to extensive transcriptional changes in maize, including up-regulation of glycolysis, the tricarboxylic acid (TCA) cycle, and defense-related genes, as well as down-regulation of photosynthetic genes.

The regulation of plant metabolism is crucial, especially when challenged by stress. Sucrose-nonfermenting1 (SNF1)-related protein kinase 1 (SnRK1) functions as a central energy regulator, consisting of a catalytic α subunit and regulatory β and γ subunits ^15^. SnRK1 regulates growth and developmental transitions under varying stresses and energy conditions by phosphorylating key transcription factors and metabolic enzymes ^16–20^. Its kinase activity is activated by the phosphorylation of a conserved threonine residue in the T-loop of the catalytic domain, mediated by upstream kinases (SnAK1/2) or via autophosphorylation ^15,21,22^. Upon activation, SnRK1 regulates a broad range of signaling and metabolic pathways by promoting catabolic processes to mobilize storage compounds, while repressing anabolic processes. The activity of SnRK1 can be inhibited by sugar phosphates or phosphatases ^23–25^. Additionally, SnRK1 modulates plant immunity against diverse plant pathogens and is targeted by pathogenic effector proteins (Hulsmans et al., 2016). For example, in pepper, SnRK1 suppressed the AvrBs1-specific hypersensitive response mediated by the effector protein AvrBsT in *X. campestris* pv. *vesicatoria* ^27^. In wheat, TaSnRK1 interacted with *Fusarium graminearum* orphan protein TaFROG to positively regulate resistance by mediating the proteasomal degradation of the cytoplasmic effector Osp24 ^28^. Overexpression of SnRK1A in rice enhanced resistance to broad-spectrum hemibiotrophic and necrotrophic pathogens ^29^. *Plasmodiophora brassicae*-specific effector PBZF1 inhibited SnRK1.1-mediated resistance in *Arabidopsis thaliana* against clubroot disease ^30^.

In this study we functionally characterized the role of Hap1-3 effectors (UMAG_02473, UMAG_00792, and UMAG_00793; Hypertrophy associated protein 1-3) in *U. maydis*, with a particular focus on Hap1. Hap effectors are required for full virulence in *U. maydis* and interact within the host cell, suggesting the potential formation of an effector complex. We demonstrate that Hap1 interacts with maize SnRK1α and the presence of Hap1 leads to the phosphorylation of SnRK1 downstream targets and cell cycle regulation, as well as the up-regulation of genes of key enzymes involved in cell cycle regulation and starch biosynthesis during infection. CRISPR-Cas9 mediated mutation of *hap1* (CR-*hap1*) reduced starch accumulation, nuclear size, and increased expression of plant defense-related genes.

## Results

### Characterization of candidate genes associated with hypertrophic tumor formation in *Ustilago maydis* effectors

To study the molecular function of HTT effectors, we selected candidate effector genes that were specifically and strongly up-regulated in hypertrophic mesophyll tumor cells in *U. maydis*-infected maize from a previously published transcriptome data set ^7^ (Table **1**). Additionally, *UMAG_00792* and *UMAG_00794,* paralogs of *UMAG_00793*, were included for further candidate screening. Genomic analysis revealed that *UMAG_00792* and *UMAG_00793*, located on chromosome 1, share an 848 bp intergenic promoter region, suggesting coordinated expression (Table **1**; Fig. **1a**). Using CRISPR-Cas9 mutagenesis in the solopathogenic *U. maydis* strain SG200 ^31^, open reading frame shift knockout mutants of candidate effector genes were generated. Mutants with reduced virulence in at least one maize line were subjected to genetic complementation with the corresponding gene under the control of its native promoter (approximately 600bp upstream). This approach identified *UMAG_02473*, *UMAG_00792*, and *UMAG_00793* as HTT-associated effector candidates, named Hap1, Hap2, and Hap3 (hypertrophic-associated proteins 1, 2, and 3), respectively (Fig. **1b**). Moreover, publicly available transcriptomic data ^32^ showed that Hap1, Hap2, and Hap3 effectors exhibited the highest gene expression at 2 dpi (Table **1**). Candidates *cda7* and *sts2* were excluded from this study, due to their established functional roles as a chitin deacetylase and a transcriptional activator, respectively ^8,33^. *UMAG_00753* was also excluded for lacking a conventional N-terminal signal peptide (SignalP 6.0; Teufel et al., 2022) and failing to exhibit secretion in an *in-planta* secretion test (Fig. **S1**).

**Fig. 1).**
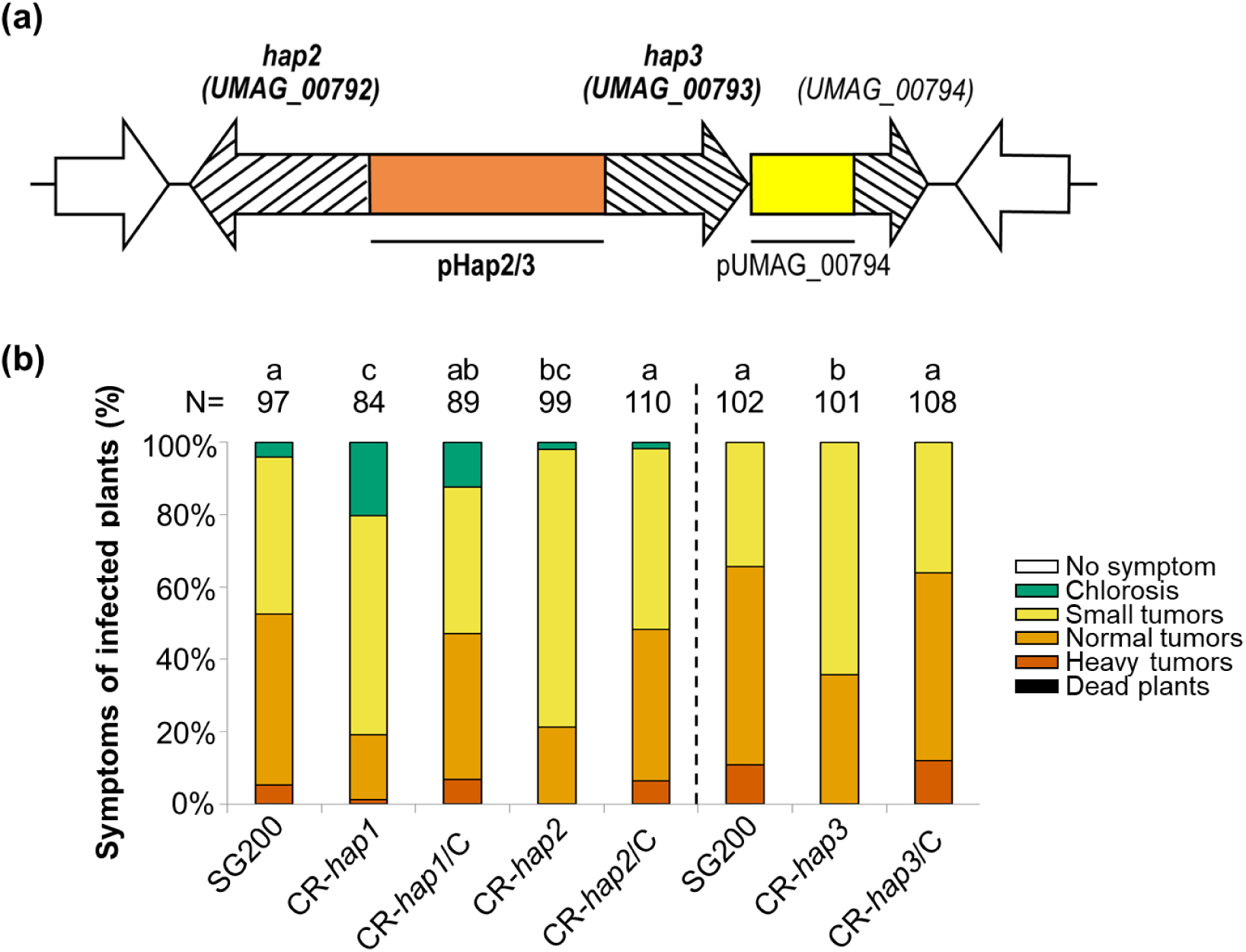
Hap effectors are required for full virulence in *U. maydis*. **(a)** Schematic representation of the organization of effector genes on chromosome 1 of *U. maydis*, showing the location of hap2, hap3, and UMAG_00794. Dashed arrows indicate three paralogous genes. The orange box represents shared promoter region of *hap2* and *hap3*. The yellow box depicts the promoter region of *UMAG_00794*. White arrows denote flanking genes that are not paralogous to the Hap effectors. **(b)** Disease symptoms in maize infected with *U. maydis* frameshift knockout mutants and complemented strains are compared to the wild-type strain SG200 at 12 dpi. The disease index is presented as the mean from three biological replicates. Statistical significance was determined using one-way ANOVA with Tukey’s HSD test (*P* < 0.05). Different letters indicate statistically significant differences among treatments.

**Table 1).**
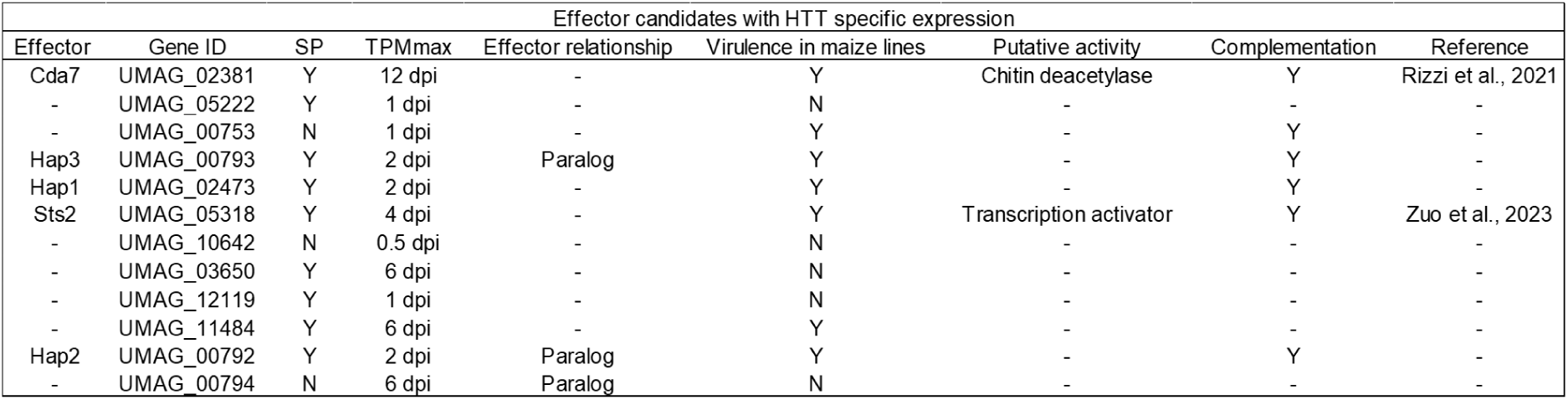
Overview of HTT-related effector candidates tested in this study. This table presents an overview of *Ustilago maydis* effector candidates related to hypertrophic tumor cells (HTT) evaluated in this study. SP, signal peptide. TPMmax, the highest Transcripts Per Million, reflecting the peak expression levels of these effectors at various plant-associated time points throughout different growth stages ^32^. Effector relationship, paralogous relationships among the HTT-related *U. maydis* effectors listed.

| Effector candidates with HTT specific expression |  |  |  |  |  |  |  |  |
| --- | --- | --- | --- | --- | --- | --- | --- | --- |
| Effector | Gene ID | SP | TPMmax | Effector relationship | Virulence in maize lines | Putative activity | Complementation | Reference |
| Cda7 | UMAG_02381 | Y | 12 dpi | - | Y | Chitin deacetylase | Y | Rizzi et al., 2021 |
| - | UMAG_05222 | Y | 1 dpi | - | N | - | - | - |
| - | UMAG_00753 | N | 1 dpi | - | Y | - | Y | - |
| Hap3 | UMAG_00793 | Y | 2 dpi | Paralog | Y | - | Y | - |
| Hap1 | UMAG_02473 | Y | 2 dpi | - | Y | - | Y | - |
| Sts2 | UMAG_05318 | Y | 4 dpi | - | Y | Transcription activator | Y | Zuo et al., 2023 |
| - | UMAG_10642 | N | 0.5 dpi | - | N | - | - | - |
| - | UMAG_03650 | Y | 6 dpi | - | N | - | - | - |
| - | UMAG_12119 | Y | 1 dpi | - | N | - | - | - |
| - | UMAG_11484 | Y | 6 dpi | - | Y | - | - | - |
| Hap2 | UMAG_00792 | Y | 2 dpi | Paralog | Y | - | Y | - |
| - | UMAG_00794 | N | 6 dpi | Paralog | N | - | - | - |

### Hap effectors interact with each other *in planta*

Hap1, Hap2, and Hap3 immunoprecipitation (IP) assays followed by mass-spectrometry were performed to identify potential effector interactors. To this end, *U. maydis* strains CR-Hap1, CR-Hap2, or CR-Hap3 strains were transformed for stable expression of the respective 2xHA-tagged *hap1*, *hap2*, or *hap3* genes under the control of the *pit2* promoter, which confers strong expression in infectious hypha ^35^. The resulting recombinant *U. maydis* strains (SG200-CR-Hap1-p*Pit2::hap1-2xHA,* SG200-CR-Hap2-p*Pit2::hap2-2xHA,* or SG200-CR-Hap3-p*Pit2::hap3-2xHA*) were used for infection of maize leaves. A *U. maydis* strain expressing HA-tagged mCherry with a signal peptide (SG200-p*Pit2::SP-mCherry-HA*) was used as a control (Fig. **2a**). Mass-spectrometry results are summarized in Table **S2-4**.

**Fig. 2).**
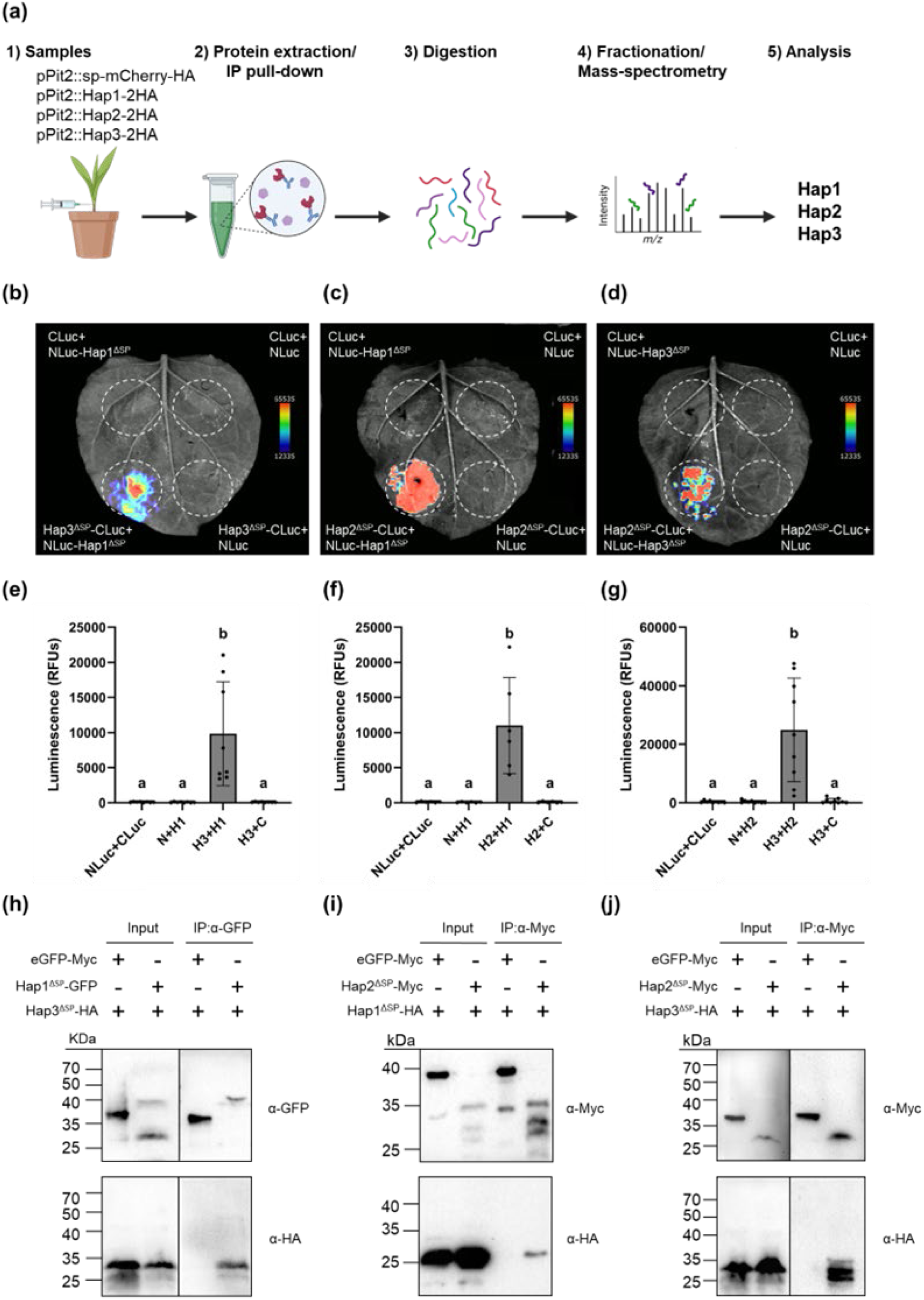
Hap effectors interact *in planta*. **(a)** Overview of pull-down/mass spectrometry (MS) workflow for identifying effector targets. (1) Seven-day-old maize seedlings were infected with *U. maydis* SG200 strains expressing HA-tagged Hap effectors and collected at 3 dpi. (2) Total proteins were extracted and immunoprecipitated using HA magnetic beads. (3) Proteins bound to the beads were digested. (4) Digested peptides were fractionated and analyzed by MS. (5) Identified spectra were mapped against the *U. maydis* genome to identify interacting effector proteins. **(b-d)** Split-luciferase complementation assay in *N. benthamiana* for confirmation of effector interaction. Hap1^ΔSP^-NLuc, Hap2^ΔSP^-NLuc, Hap3^ΔSP^-NLuc, or empty-NLuc were co-expressed with Hap1^ΔSP^-CLuc, Hap2^ΔSP^-CLuc, Hap3^ΔSP^-CLuc, or empty-CLuc. Luminescence was detected using Bio-Rad ChemiDoc™ and pseudo-fluorescence applied for enhanced visualization. A representative image from three independent replicates is shown. **(e-g)** Quantification of luciferase activity, shown as relative luciferase units (RFU), from the data presented in Fig. 3C. Error bars represent standard deviation; *n* = 3 biological replicates. Statistical analysis was performed using one-way ANOVA with Tukey’s HSD test (*P* < 0.05). Different letters indicate statistically significant differences among treatments. **(h-j)** Co-immunoprecipitation (Co-IP) assay in *N. benthamiana*. p2x35S-Hap1^ΔSP^-GFP or p2x35S-Hap2^ΔSP^-4xMyc were co-expressed with p2x35S-Hap1^ΔSP^-6xHA or p2x35S-Hap3^ΔSP^-6xHA, and controls with p2x35S-Hap1^ΔSP^-6xHA with p2x35S-GFP or p2x35S-GFP-4xMyc. Proteins pulled-down with GFP or Myc magnetic beads were detected using anti-HA, anti-GFP, or anti-Myc antibodies. Expected protein sizes: Hap1^ΔSP^-GFP=40.05 kDa; Hap2^ΔSP^-4xMyc=23.11 kDa; Hap1^ΔSP^-6xHA=20.07 kDa; Hap3^ΔSP^-6xHA=23.22 kDa; GFP=27 kDa; GFP-4xMyc=31.8 kDa. N: NLuc; C: CLuc; H1: Hap1; H2: Hap2; H3: Hap3

Surprisingly, a significant number of the detected Hap-interactors were found among the 467 predicted secreted effector proteins of *U. maydis* (Kämper et al., 2006; Lanver et al., 2017). An enrichment of 32, 4, and 13 proteins was found for the Hap1, Hap2, and Hap3 samples, respectively. Subsequent analyses using EffectorP_Fungi 3.0, SignalP 6.0, TPMmax, InterproScan identified secreted cytoplasmic effectors with the highest expression at 2 dpi ^32,34,37,38^. Pull-down assays detected five cytoplasmic and dual-localized apoplastic/cytoplasmic effectors in Hap1, three cytoplasmic and dual-localized cytoplasmic/apoplastic effectors in Hap2, and two cytoplasmic effectors were detected in Hap3 (Data **S2-4**). Notably, among these putative interactors, only Hap1, Hap2, and Hap3 proteins co-precipitated with each other, indicating *in planta* interactions amongst Hap1, Hap2, and Hap3 (Fig. **S2a-c**).

To verify the interactions amongst Hap effectors, a split luciferase complementation assay was performed. In this assay, nLuc-Hap1^ΔSP^, -Hap2^ΔSP^, or -Hap3 ^ΔSP^ was transiently co-expressed with Hap1^ΔSP^-, Hap2^ΔSP^-, or Hap3 ^ΔSP^-cLuc in *Nicotiana benthamiana* using *Agrobacterium*-mediated transformation. Luminescence signals were only detected when Hap effectors were co-expressed, while no visible signal was observed when Hap effectors were co-expressed with empty-cLuc or -nLuc controls (Fig. **2b-g**). To further validate the interaction amongst Hap effectors, co-immunoprecipitation (co-IP) assays were performed using α-Myc or α-GFP magnetic beads in *N. benthamiana*. *Agrobacterium* strain carrying Hap1^ΔSP^-GFP or Hap2 ^ΔSP^-4xMyc was co-infiltrated with Hap3^ΔSP^-6xHA or Hap1^ΔSP^-6xHA. As a negative control, eGFP or GFP-4xMyc was co-infiltrated with Hap3^ΔSP^-6xHA or Hap1^ΔSP^-6xHA. Hap1^ΔSP^-GFP or Hap2^ΔSP^-4xMyc was co-immunoprecipitated with Hap1^ΔSP^-6xHA or Hap3^ΔSP^-6xHA, but not in either of eGFP or GFP-4xMyc controls (Fig. **2h-j**). Collectively, these results show that Hap effectors interact with each other *in vivo*.

Given the overlapping, HTT-specific expression patterns of Hap1, Hap2, and Hap3 at 2dpi (Table 1), as well as their interaction *in planta*, we hypothesized that these effectors might have additive or cooperative effects on virulence. To assess their collective contribution, we generated triple frameshift knockout mutants for *hap1/2/3* using CRISPR-Cas9 mutagenesis. Interestingly, the virulence level of the resulting triple frameshift knockout mutant was similar to that of the single *hap1* frameshift knockout mutant, suggesting that while *hap2* and *hap3* contribute to the effector complex, *hap1* plays a dominant role as the HTT-associated effector driving *U. maydis* virulence (Fig. **S3**).

### Hap effectors influence Hap1-ZmSnRK1α interaction and metabolic processes in maize

To elucidate maize proteins targeted by Hap1, identified spectra from the above-described IP-MS/MS analyses were mapped to the maize proteome (Fig. **2a**, Mass-spectrometry result is summarized in Table **S5**). Top 150 proteins detected in Hap1-expressing samples were analyzed with Gene Ontology (GO) enrichment using PLAZA 5.0 and grouped into higher hierarchical terms by REVIGO, revealing that Hap1 targets maize proteins involved in protein phosphorylation and serine-threonine kinase activity, particularly isoforms of the SnRK1α subunit (Fig. **S4a and b**).

To confirm an interaction between Hap1 and ZmSnRK1α, a split-luciferase complementation assay was conducted. In this assay, Hap1^ΔSP^-cLuc was transiently co-expressed with nLuc-ZmSnRK1α1, -ZmSnRK1α2, or -ZmSnRK1α3 in *N. benthamiana* using *Agrobacterium*-mediated transformation. Empty-nLuc or -cLuc were used as a control. Luminescence signals were detected exclusively when Hap1^ΔSP^-cLuc was co-expressed with nLuc-ZmSnRK1α2 or nLuc-ZmSnRK1α3, while no visible signal was observed with nLuc-ZmSnRK1α1, empty-cLuc, or -nLuc controls (Fig. **3a and b**). To further validate the interaction between Hap1 with ZmSnRK1α, co-IP assays in *N. benthamiana* were conducted. *Agrobacterium* strain carrying Hap1^ΔSP^-6xHA were co-infiltrated with ZmSnRK1α1-4xMyc, ZmSnRK1α2-4xMyc, or ZmSnRK1α3-4xMyc. As a negative control, GFP-4xMyc was co-infiltrated with Hap1^ΔSP^-6xHA. Agroinfiltrated leaves of ZmSnRK1α1, ZmSnRK1α2, or ZmSnRK1α3 were co-immunoprecipitated by α-Myc immunoprecipitation of Hap1^ΔSP^-6xHA, but not in GFP-4xMyc control (Fig. **3c**). These results show that Hap1 interacts with catalytic subunits of ZmSnRK1.

**Fig. 3).**
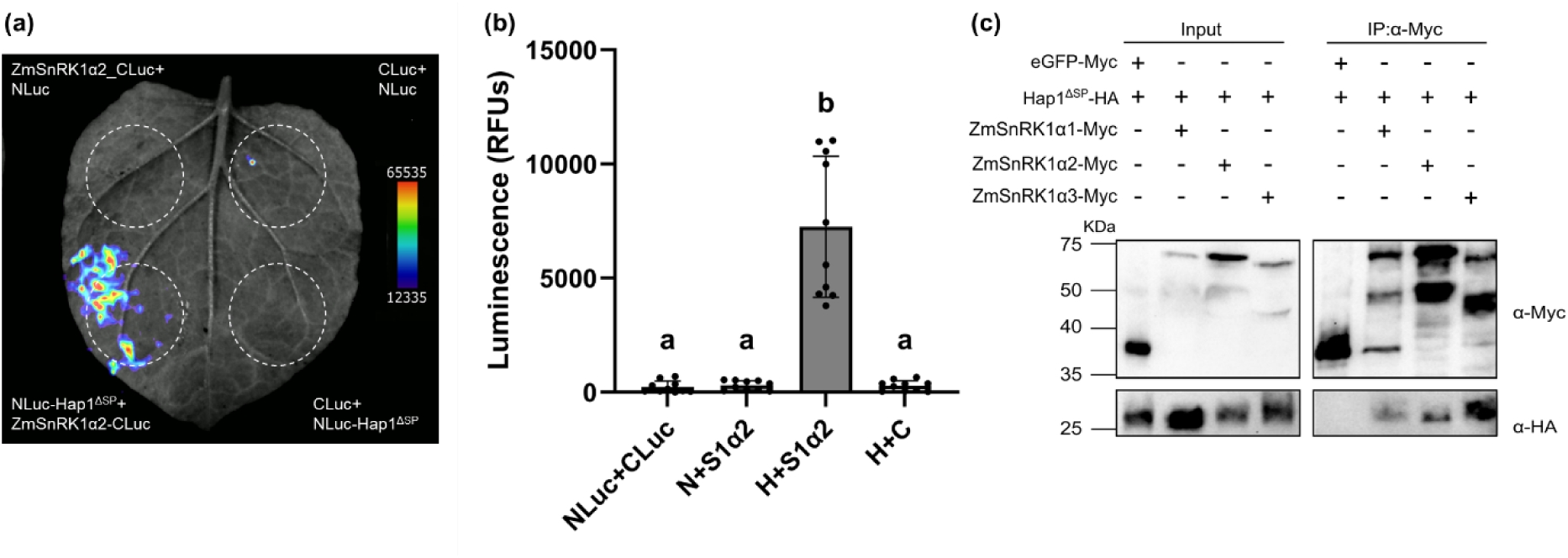
Hap1 interacts with maize SnRK1α. **(a)** Split-luciferase complementation assay in *N. benthamiana*. Hap1^ΔSP^-NLuc or empty-NLuc were co-expressed with ZmSnRK1α2-CLuc, ZmSnRK1α3-CLuc, or empty-CLuc. Luminescence was detected using Bio-Rad ChemiDoc™ and pseudo-fluorescence applied for enhanced visualization. Representative image of ZmSnRK1α2 from three independent biological replicates and ZmSnRK1α3 from two biological independent replicates. **(b)** Quantification of luciferase activity, shown as relative luciferase units (RFUs), from the data presented in Fig. 4A. Error bars represent standard deviation; *n* = 3 biological replicates. Statistical analysis was performed using one-way ANOVA with Tukey’s HSD test (*P* < 0.05). Different letters indicate statistically significant differences among treatments. N: NLuc; C: CLuc; H: Hap1; S1α2: ZmSnRK1α2. **(c)** Co-immunoprecipitation (Co-IP) assay in *N. benthamiana*. p2x35S-Hap1^ΔSP^-6xHA was co-expressed with p2x35S-ZmSnRK1α1-4xMyc, p2x35S-ZmSnRK1α2-4xMyc, or p2x35S-ZmSnRK1α3-4xMyc and controls with p2x35S-Hap1^ΔSP^-6xHA with p2x35S-GFP-4xMyc. Proteins pulled down with Myc magnetic beads were detected using anti-HA or anti-Myc antibodies. Expected protein sizes: Hap1^ΔSP^-6xHA=20.07kDa; ZmSnRK1α1-4xMyc=64.23kDa; ZmSnRK1α2-4xMyc=63.27kDa; ZmSnRK1α3-4xMyc=62.96kDa; GFP-4xMyc=31.8 kDa.

Building on our findings from interaction studies of Hap effectors and Hap1-ZmSnRK1, we hypothesized that maintaining Hap effectors as a complex is important for orchestrating a regulatory cascade and mediating interactions between Hap1 and host proteins. To test this hypothesis, we generated single frameshift knockout mutants for *hap2* and *hap3* (SG200-CR-Hap1-p*Pit2::Hap1-2xHA-CR-hap2* and SG200-CR-Hap1-p*Pit2::Hap1-2xHA-CR-hap3*) as well as a double frameshift mutant for both hap2 and hap3 (SG200-CR-Hap1-p*Pit2::Hap1-2xHA-CR-hap2-3*) using CRISPR-Cas9 mutagenesis in SG200-CR-Hap1-p*Pit2::Hap1-2xHA* background strain. Maize seedlings were subsequently infected with the respective mutants, and SG200-pPit2::SP-mCherry-HA strain was used as a control for co-IP followed by MS analysis (Fig. **4a**, mass-spectrometry results are summarized in Table **S6-8)**. Proteins exclusively detected in frameshift mutants were subjected to GO enrichment analysis using PLAZA 5.0 and PPI analysis by STRING database. GO and PPI results revealed significant association in ‘cellular component organization or biogenesis’ and ‘Protein phosphatase activity’ in Hap1-CR-Hap2; ‘protein modification’, ‘brassinosteroid-mediated signaling’, ‘starch metabolic process’ in Hap1-CR-Hap3; ‘ER-Golgi transport,’ ‘calcium channel activity,’ and ‘hydrolase activity’ in the Hap1-CR-Hap2/3 double frameshift mutant (Fig. **4b**, **S5a-c**).

**Fig. 4).**
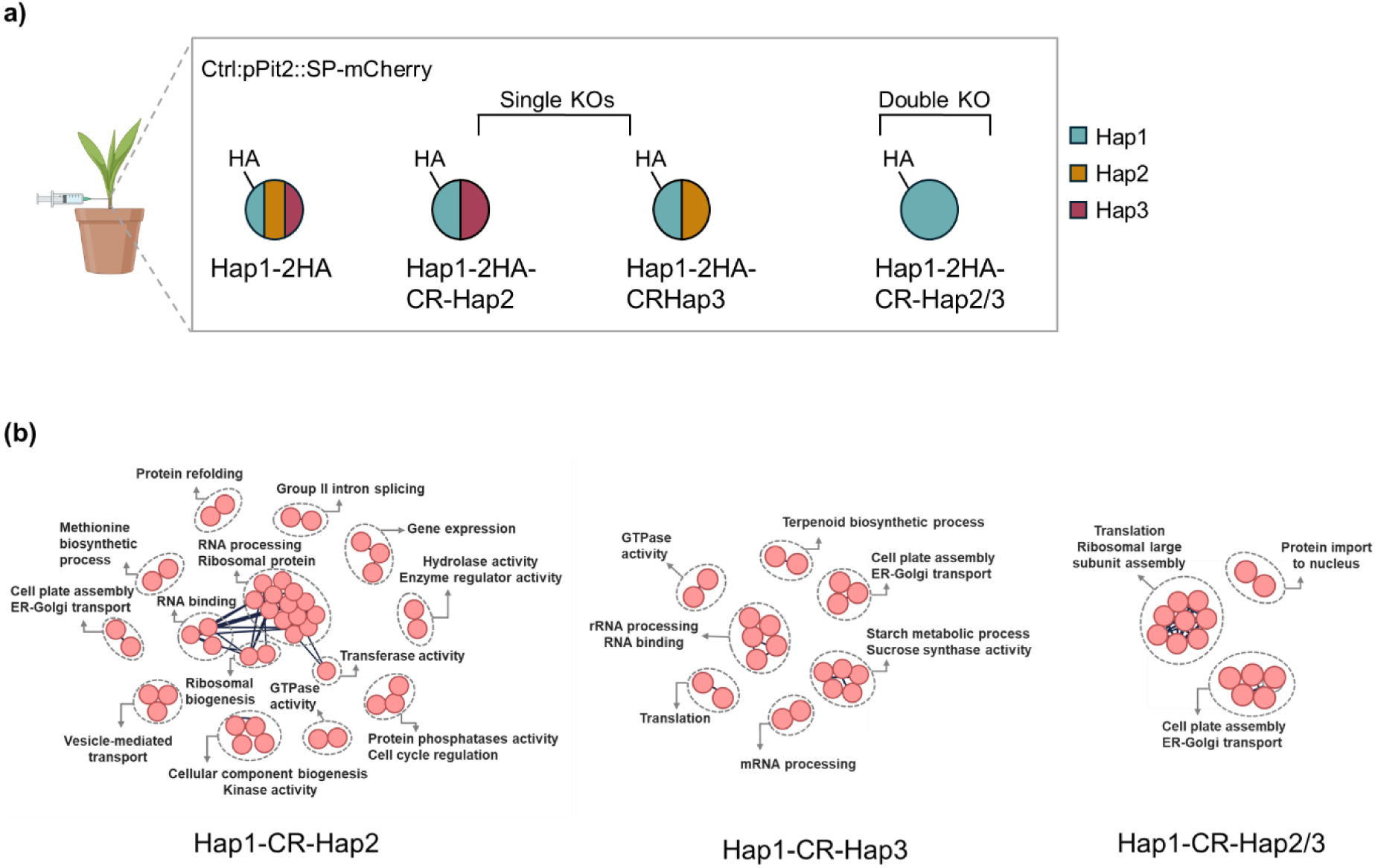
Alterations in Hap effectors modify the proteomic profile of Hap1. **(a)** IP/MS was conducted for host target identification of Hap1-CR-Hap2, Hap1-CR-Hap3, and Hap1-CR-Hap2/3. 7-day-old maize seedlings were infected with *U. maydis* SG200 strains expressing SG200-CR-Hap1-p*Pit2::Hap1-2xHA-CR-Hap2,* SG200-CR-Hap1-p*Pit2::Hap1-2 xHA-CR-Hap3*, SG200-CR-Hap1-p*Pit2::Hap1-2xHA-CR-Hap2-3,* or SG200-p*Pit2::SP-mCherry-HA*. Blue represents Hap1 protein, yellow represents Hap2 protein, and red represents Hap3 protein. **(b)** The PPI network was constructed using the STRING database, with nodes demonstrating individual proteins and edges representing predicted functional associations and interactions. Default settings with a high confidence threshold (0.7) were applied, and the network was clustered using K-means clustering. The connections shown are manually curated to highlight important interactions.

In light of observed analysis above, we focused on kinases and SnRK1 substrates, given that SnRK1α is the primary host target(s) of Hap1. Interestingly, all frameshift knockout mutants interacted with phosphatases and trehalose-6-phosphate synthases, known SnRK1 substrates. Specifically, Hap1-CR-Hap2 interacted with poorly characterized proteins, such as protein phosphatase 2C (PP2C), a regulator of the signal transduction pathway involved in modulating receptor-like kinases and abscisic acid signaling ^39^, and phosphotyrosyl phosphatase activator (PTPA) which activates protein phosphatase 2A (PP2A). Moreover, both Hap1-CR-Hap2 and Hap1-CR-Hap2/3 interacted with PPP2CB, the catalytic subunit of PP2A that negatively regulates cell cycle progression by controlling the timing and coordination of cell division ^40^. In contrast, Hap1-CR-Hap3 interacted with enzymes involved in starch biosynthesis, as well as kinases and phosphatases associated with the brassinosteroid signaling pathway. Interestingly, both Hap1-CR-Hap3 and Hap1-CR-Hap2/3 mutants interacted with SnRK2, a kinase involved in stress and abscisic acid (ABA)-mediated signaling pathways, as well as fructose-2,6-biphosphatases, which are important in regulating glycolysis and gluconeogenesis (Data **S5**).

### Hap1 contributes to starch reprogramming and endoreduplication in maize HTT cells

To investigate the host transcriptional response to *U. maydis* infection, RNA sequencing was performed on maize seedlings infected with mock, SG200, or CR-*hap1* at 3 dpi, using three independent biological replicates. Comparisons between *U. maydis*-infected (SG200 or CR-*hap1*) and mock-infected samples (i.e., S vs. M and CR-H vs. M) revealed 12,492 and 10,603 differentially expressed genes (DEGs), respectively (Data **S6 and 7**). Specifically, in S vs. M comparisons, 7,676 up-regulated and 4,816 down-regulated were identified, while in CR-H vs. M, 6,795 up-regulated and 3,808 down-regulated were identified (Fig. **S6a**). Of these, 5,974 up-regulated and 3,072 down-regulated DEGs were common in both comparisons (Fig. **S6b and c**). GO analysis of shared up-regulated DEGs revealed enrichment in biological processes related to ‘carbohydrate derivative metabolic process’, and ‘carbohydrate catabolic process’. These results indicate that *U. maydis* influences host cellular activities related to complex molecule synthesis and carbohydrate metabolism.

To gain further insight into the impact of *hap1* on host gene expression during *U. maydis* infection, we performed a pairwise comparison of CR-*hap1* and SG200 treatments (i.e., CR-H vs. S), identifying 908 up-regulated and 1,431 down-regulated DEGs out of 2,339 genes (Fig. **5a**, Data **S8**). GO enrichment analysis of the up-regulated DEGs in CR-*hap1* revealed biological processes related to defense responses such as ‘cell surface receptor signaling pathway’ and ‘response to oomycetes’ (Fig. **S7a**). Among these, six WRKY transcription factors were found to be up-regulated, including *WRKY82,* which regulates the phenolic pathway in maize ^41^. *WRKY17* was induced upon *U. maydis* infection and interacts with the SWEET4b transporter in yeast-one-hyrid assay ^42^. In Arabidopsis, the *WRKY17* homolog activates SA-dependent defense genes, while repressing JA-regulated genes ^43^. Moreover, cell surface receptor signaling pathways, such as wall-associated kinases and lectin domain-containing proteins, were also up-regulated, which are important for plant immunity against pathogenic attack (Fig. **S7b**, Data **S9**). Conversely, up-regulated DEGs in SG200 were linked to cell cycles such as ‘regulation of cell cycle’, ‘cell division’, and ‘DNA replication’ (Fig. **5b**). Notably, 104 cell cycle-related genes were up-regulated in the presence of *hap1*, especially those involved in G1/S phase transitions, such as *WEE1* kinase, *SMR1* (siamese-related1), *SMR3* (siamese-related3), three D-type cyclins, and pre-replication complex (pre-RC) components, such as minichromosome maintenance 3-7 (*MCM3-7*) Origin recognition complex 5-6 (*ORC5-6*), cell division cycle 6 (*CDC6*), Cdc10-dependent transcript 1 (*CDT1*), as well as replication protein A 1-3 (*RPA1-3*), which binds to single-stranded DNA during replication, DNA polymerase alpha subunit 2 (*POLA2*), which anchors the polymerase alpha-primase complex to the replication fork, and two E2F-DP coding genes required for the onset of the S phase ^44–56^ (Fig. **5c**, Data **S10**).

**Fig. 5).**
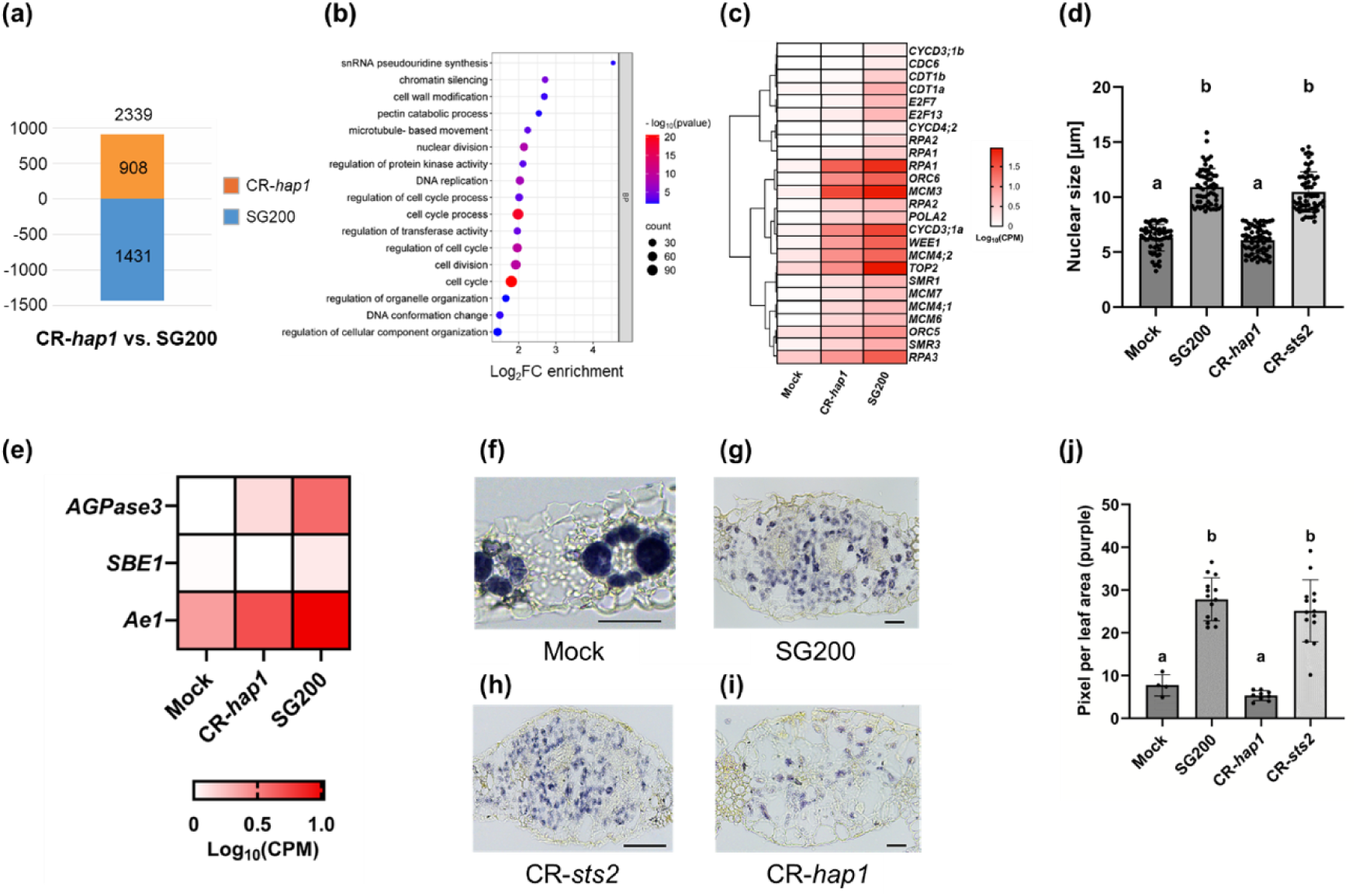
The presence of Hap1 is required for starch accumulation and endoreduplication in *U. maydis* infected HTT cells. **(a)** Pairwise bar graph depicting the number of up- and down-regulated DEGs in CR-*hap1* vs. SG200 **(b)** GO enrichment analysis of DEGs up-regulated in SG200. **(c)** Heatmap of DEGs related to cell cycle processes, specifically involved in G-S1 phase transition. DEGs = Log_2_FC>1 or <-1 and FDR <0.05. **(d)** Quantification of histological cross-sections of leaf tissues infected with mock or *U. maydis* (SG200, CR-*hap1,* CR-*sts2*) stained with propodium iodied at 6 dpi. Quantification was performed from three independent biological replicates, each with > 55 nuclei per sample. **(e)** Heatmap of DEGs related to key enzymes in starch biosynthesis in SG200-infected samples. DEGs = Log_2_FC>1 or <-1 and FDR <0.05. Histological cross-sections of leaf tissues infected with mock or *U. maydis* (SG200, CR-*hap1,* CR-*sts2*) stained with Lugol iodide (IKI) at 6 dpi: **(f)** Mock, **(g)** SG200, **(h)** CR-*hap1*, **(i)** CR-*sts2*. Representative image of SG200, CR-*hap1*, and CR-*sts2* from three independent biological replicates and mock from two independent biological replicates. Scale bars = 50 μm. **(j)** Quantification of starch staining from Fig.7f-i. Statistical analysis for panels d and j were performed used one-way ANOVA with Tukey’s HSD test (*P* < 0.05). Different letters indicate statistically significant differences among treatments.

To further explore the potential role of Hap1 in endoreduplication and starch synthesis in *U. maydis*-infected mesophyll HTT cells, maize seedlings were infected with mock or *Ustilago* (SG200, CR-*hap1*, and CR-*sts2*) strains. *U. maydis* strain CR-*sts2*, in which the transcription activator effector Sts2 that promotes *de novo* cell division in bundle sheath tumor cells (HPT) is disrupted ^8^, was used as a positive control. Infected plant tissues were collected at 6 dpi, embedded in paraplast, and thin-sectioned for analysis. Endoreduplication was assessed by staining the leaf sections with propidium iodide (PI) to measure nuclear size of mesophyll cells. Interestingly, CR-*hap1* infected tissues exhibited nuclear sizes similar to mock, but approximately two times smaller than those infected with SG200 and CR-*sts2* (Fig. **5d**).

Cell cycle progression is controlled by several factors and one such factor in plants is sugar signaling ^57^. Thus, DEGs related to sugar and starch metabolism were investigated. Notably, we found increased gene expression levels of key starch biosynthetic enzymes, such as *ADP-glucose pyrophosphorylase 3* (*AGPase 3*), *starch-branching enzyme I* (*SBE I*), and *amylose extender 1* (*Ae1*) in maize leaves infected with SG200 (Fig. **5e**, Data **S11**). To examine starch distribution, tissue sections were stained with Lugol’s iodine solution (IKI). In mock-infected tissues, starch accumulation was confined to the bundle sheath, whereas SG200-infected tissues showed dispersed and increased starch accumulation in mesophyll cells, consistent with previous studies ^7^ (Fig. **5f-g** and **j**). Notably, CR-*hap1*-infected tissues displayed significantly reduced starch accumulation in mesophyll compared to SG200 (Fig. **5h** and **j**). CR-*sts2* sections showed no significant difference in starch accumulation compared to SG200, indicating that reduced starch accumulation is a specific characteristic of CR-*hap1* (Fig. **5i** and **j**).

### Phosphoproteomic analysis of maize upon *Ustilago maydis* infection

To investigate the impact of Hap1 on SnRK1-dependent phosphorylation, a quantitative phosphoproteomic analysis using mock-infected and *Ustilago*-infected (SG200 and CR-Hap1) was performed (Fig. **6a**). Infected maize seedlings were collected at 3 dpi in three independent biological replicates. To identify proteins specifically enriched due to phosphorylation rather than general protein abundance, a Venn analysis comparing the phosphoproteome and total proteome of *Ustilago*-infected (SG200 or CR-Hap1) samples with mock-infected sample, using a cut-off of Log_2_FC≥1 or ≤-1, was performed. In the phosphoproteome analysis of SG200-infected samples compared to mock (S vs. M), 3,618 phosphopeptides derived from 2,337 proteins were increased and 180 phosphopeptides derived from 148 proteins were decreased. In the total proteome, S vs M analysis revealed 1,534 enriched and 745 depleted proteins (Fig. **S8a and b**). Integration of these datasets using Venn analysis resulted in 1,983 increased phosphorylated proteins and 115 decreased phosphorylated proteins (Fig. **S8a**). Similarly, in Hap1-infected compared to mock-infected (CR-H vs. M), the phosphoproteome revealed 1,480 increased phosphopeptides from 1,205 proteins and 117 decreased phosphopeptides from 94 proteins. In the total proteome of CR-H vs. M, 1,086 enriched and 567 depleted proteins have been identified, respectively (Fig. **S8c and d**). Integrating these datasets resulted in 1,028 increased phosphorylated proteins and 64 decreased phosphorylated proteins (Fig. **S8c**). Interestingly, the number of phosphorylated proteins increased in S vs. M was approximately twice that in CR-H vs. M, suggesting that the absence of Hap1 may hinder interactions with targets involved in post-translational modifications. Total proteomic and phosphoproteomic data are summarized in Table **S9 and 10**.

**Fig. 6).**
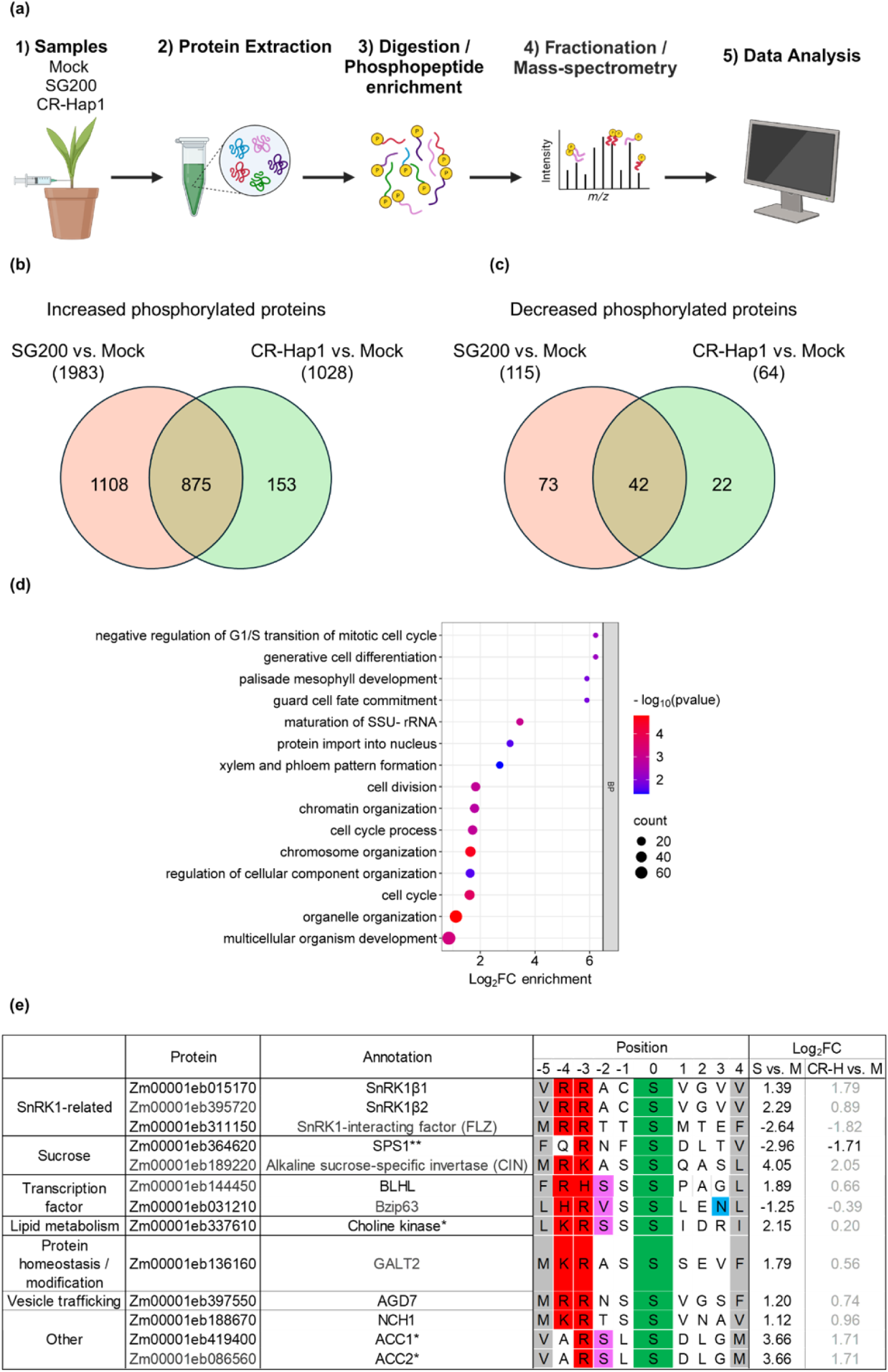
Phosphoproteomic analysis of maize infected with CR-Hap1. **(a)** Schematic overview of the phosphoproteomics experiment: (1) seven-day-old maize seedlings were infected with mock, SG200, or SG200_CR_Hap1 and collected at 3 dpi. (2) Total maize proteins were extracted and separated into total proteome and phosphoproteome fractions. (3) Protein peptides were phosphoenriched using titanium dioxide. (4) Peptides were fractionated and analyzed by mass spectrometry. (5) Identified spectra were mapped to the *Z. mays* genome to identify phosphorylation sites. **(b)** Venn diagram showing uniquely increased phosphorylated proteins in SG200 and CR-Hap1 compared to mock. **(c)** Venn diagram showing uniquely decreased phosphorylated proteins in SG200 and CR-Hap1 compared to mock. **(d)** GO enrichment analysis of proteins with increased phosphorylation in SG200 vs. CR-Hap1. Significance is indicated by −Log10(*P*-value), with color shading from red (high significance) to blue (low significance). **(e)** Overview of the SnRK1-dependent consensus motif at P-5 and P+4 relative to phosphorylated Ser/Thr residues. The color scheme follows the MEME Suite’s coding for amino acids. Amino acid matching the known human AMPK consensus phosphorylation sequence is colored in pink and blue at P-2 and P+3 positions, respectively. Log_2_FC>1 or <-1 and *p* <0.05.

To differentiate Hap1-dependent from Hap1-independent phosphorylation, we compared uniquely increased or decreased phosphorylated proteins in S vs. M and CR-H vs. M. The analysis revealed 1,108 and 73 proteins with increased and decreased phosphopeptides in S vs. M. In CR-H vs. M, 153 and 22 increased or decreased phosphorylated proteins were identified. Surprisingly, 85% (875) of phosphorylated proteins in CR-H vs. M overlapped with those in S vs. M, indicating that SG200 and CR-Hap1 share congruent regulatory responses and signaling events, with the absence of Hap1 having minimal impact (Fig. **6b and c**). GO enrichment analysis of increased proteins containing phosphopeptides in S vs. M revealed biological processes of ‘negative regulation of G1/S transition of mitotic cell cycle, ‘peptidyl-serine modification’, ‘negative regulation of metabolic process, ‘protein autophosphorylation’, and ‘regulation of gene expression’ (Data **S12**). Analysis of increased phosphorylated proteins in CR-H vs. M revealed ‘negative regulation of protein-containing complex disassembly’ and ‘response to stimulus’ (Data **S13**). To further study Hap1-dependent phosphorylation changes in maize upon infection, a comparison between SG200 and CR-Hap1 (S vs. CR-H) was performed, resulting in 332 increased and 15 decreased phosphorylated proteins. GO enrichment analysis of the biological processes of increased phosphoproteins in SG200-infected leaf tissue revealed ‘negative regulation of G1/S transition of mitotic cell cycle, ‘cell division’ and ‘palisade mesophyll development (Fig. **6d**). In SG200-infected tissue, increased phosphorylation was observed in several key proteins, including two retinoblastoma-related (RBR) proteins (RBR1 and RBR3), which function as tumor suppressors by negatively regulating the cell cycle ^58–60^.

The data shown so far indicates that Hap1 facilitates endoreduplication of mesophyll HTT cells by disrupting cell cycle regulation, potentially contingent upon carbon resource availability in the tumor. Consequently, we focused our search on the known SnRK1 substrates and their consensus motif (phiXXXX**S/T**XXXphi), characterized by 10-amino acid sequence in which phi represents hydrophobic residues (M, L, V, I, or F) at positions P-5 and P+4, and a basic residue at positions P-3 and P-4 relative to the serine/threonine residue ^18^. Additionally, valine or serine at position P−2 and aspartic acid or asparagine at position P+3 of the human AMPK consensus motif was considered as selection criteria ^61^. Using the Find Individual Motif Occurrences (FIMO) database, the occurrence of SnRK1 consensus sequence on 1,108 increased and 73 decreased unique to S vs. M, and 153 increased and 22 decreased unique to CR-H vs. M were searched. In addition, increased and decreased phosphorylated proteins from S vs. CR-H were scanned to search for Hap1-dependent and independent phosphorylation. A total of 67 motifs matching the known SnRK1 consensus sequence among increased phosphorylated proteins and 6 motifs of the decreased phosphorylated proteins were found in S vs. M, while in CR-H vs. M, 13 increased and 1 decreased motif of phosphorylated proteins were found. Finally, cross-referencing with phosphoproteomic data resulted in 13 proteins perfectly matching the SnRK1 consensus sequence (Fig. **6e**). Relaxing the stringency criteria to include motifs with only one of the hydrophobic residues or only hydrophobic residues at (P−5 or P+4) found 25 proteins matching the SnRK1 consensus sequence (Data **S14**). However, these motifs should be interpreted with caution due to their similarity to the phosphorylation motifs of calcium-dependent protein kinases (CDPKs) ^18,62,63^. Remarkably, in our phosphoproteomic analysis, we found that SnRK1α1, β1 and β2, and γ are highly phosphorylated in S vs. M, but not in CR-H vs. M. This suggests that the presence of CR-Hap1 is indispensable for SnRK1 complex integrity and formation. However, the phosphopeptide identified in SnRK1α1 did not align with the conventional SnRK1 T-loop phosphorylation site. Analysis of SnRK1 consensus sequences across different conditions revealed several key enzymes involved in primary metabolism. In S vs. M, increased phosphorylation was observed in sucrose synthase 2 (Susy 2), Susy 7, and Alkaline sucrose-specific invertase (CINV), while decreased phosphorylation was observed in transcription factor bZIP63 and poorly characterized phosphatase C (PP2C) and FCS-like zink finger (FLZ). In S vs. CR-H, choline kinase, involved in lipid metabolism showed increased phosphorylation, while decreased phosphorylation was observed in sucrose phosphate synthase 1 (SPS1), an enzyme involved in sucrose biosynthesis.

## Discussion

In this study, we investigated the roles of *U. maydis* effectors being specifically activated in hypertrophic tumor cells. We identified the effector proteins Hap1, Hap2, and Hap3 as virulence factors, with Hap1 holding a key role in the induction of hypertrophy in maize leaves. Our data shows that within the host tissue, Hap effectors interact with each other, and Hap1 interacts with ZmSnRK1, a major regulator of energy and stress responses. Notably, disruption of one or both of the Hap2 and Hap3 altered Hap1’s interaction partners, such as uncharacterized PP2C, PP2A, and ZmSnRK2. The PP2C phosphatases such as ABI1, and PP2CA are negative regulators that repress ABA/SnRK2 signaling, dephosphorylating SnRK1 T-loop in *Arabidopsis* ^25^. The observed interactions suggest that Hap effectors form an effector complex and function in a compensatory manner, forming a regulatory cascade that manipulates host signaling pathways for the benefit of the pathogen.

### Hap1 modulates SnRK1-mediated energy signaling

SnRK1 activity is regulated by sugar metabolites, including glucose-1-phosphate (G1P), G6P, T6P, and ribose 5-phosphate (R5P), with T6P being a potent inhibitor ^23,24,64^. In *U. maydis*-induced leaf tumors, T6P, G1P, and G6P accumulated and showed highest the gene expression from 2 dpi-4 dpi, coinciding with the peak expression of Hap1 ^12,32^. Starch, T6P, and SnRK1 exhibit a regulatory interconnection, with T6P functioning as a signaling molecule that modulates AGPase activity via redox activation thereby influencing starch synthesis (Kolbe et al., 2005; Tiessen et al., 2003; Lunn et al., 2006). Furthermore, increasing sucrose levels proportionally raised T6P levels, suggesting that T6P acts as a signaling molecule between cytosol and the chloroplast without changing the rate of photosynthesis ^65,66^. The role of SnRK1 differs between source and sink tissues. Overexpression of AKIN10 in potato increased starch accumulation but led to reduced starch levels in *Arabidopsis* seedlings ^67,68^.

Hap1 was found to be crucial for promoting starch redistribution within the HTT cell and up-regulating genes of key starch biosynthesis enzymes, including Zm*AGPase3*, Zm*SBEI*, and Zm*Ae1*. AGPase3, the first enzyme in starch biosynthesis, converts glucose-1-phosphate to ADP-glucose, which is essential for synthesizing amylose and amylopectin. Starch-branching enzymes (SBEI and SBEII) further modulate structure and functional properties of starch ^69^. *Ae1* encodes SBEIIb, which is important for the branching of amylopectin, a major component of starch in maize ^70,71^. These up-regulation of genes transform HTT cells into nutrient sinks that favor fungal colonization and align with reports showing that *U. maydis* modulate carbohydrate metabolism, inducing starch accumulation and reallocation in infected tumorous mesophyll cells ^7,72^. Similar mechanisms were seen in other plant pathogens, such as *Spongospora subterranea f. sp. subterranean*, a soilborne protist, that manipulates starch-degrading enzymes in infected roots to promote sporosorus development and up-regulate genes involved in starch synthesis within the root galls ^73^.

### Phosphorylation of key enzymes in carbon metabolism

SnRK1 plays a central role in maintaining energy homeostasis by phosphorylating enzymes like SPS and Susy. In the active state, SnRK1 phosphorylates SPS, reducing sucrose synthesis to conserve energy by limiting energy-intensive processes ^74,75^. We found SPS phosphorylation being increased at the conserved S^162^ residue in mock samples (S vs. M, CR-H vs. M) and CR-H (CR-H vs. S), indicating that SPS is inactivated in uninfected tissues with normal photosynthetic functions to prevent excessive sucrose accumulation that could disrupt cellular balance. In contrast, SPS was dephosphorylated and remained active in *U. maydis*-infected samples (SG200 and *CR-*Hap1) and this observed phosphorylation was greater in SG200-infected samples. This suggests that the presence of Hap1 in *U. maydis* maintains SnRK1 active regardless of cellular sugar level in the tumorous tissues to activate SPS, contributing to the formation of nutrient-rich sink tissues.

Susy, another target of SnRK1, cleaves sucrose into UDP-glucose and ADPG, a precursor of glycolysis, starch, and cellulose biosynthesis ^76,77^. In phosphoproteomic analysis, Susy7 showed increased phosphorylation in S vs. M, while Susy2 exhibited increased phosphorylation in both S vs M and dH vs M comparisons, with greater increase in S vs. M. Previous studies identified SnRK1 phosphorylates ZmSusy1 at residues S^15^ and S^170^, indirectly regulating sucrose degradation and starch biosynthesis ^78,79^. Our data demonstrated that Susy7 is increased in phosphorylation at residues S^12^ and S^16^, and Susy2 at S^11^, which showed conservation with the known sucrose phosphorylation sites that have shown catalytic activities and are potential targets of SnRK1 ^79–81^. In crops, SnRK1 primarily phosphorylates Susy in sink tissues to facilitate starch accumulation ^82^. Nevertheless, we observed increased phosphorylation of Susy2 and Susy7 in *U. maydis*-infected samples in the presence of Hap1, suggesting that infected tumor cells function as sink tissues.

### The role of Hap1 in maize cell cycle progression

Plant cell expansion is linked to internal sugar availability derived from photosynthesis, with lower internal sugar concentrations associated with chloroplast differentiation and starch accumulation ^83,84^. Sucrose influences cell cycle progression, particularly the G1 to S phase transition, by upregulating genes of cyclins such as *CYCD3;1* and components of the pre-replication complex (pre-RC), which license DNA replication during G1 ^85–88^. In maize, D-type cyclins are predominantly expressed in root meristems and are less abundant in differentiating tissues; however, Zm*CycD3;1a* showed high transcript levels in the mesocotyl and leaf, suggesting a role of D-type cyclins in endoreduplication ^45^. Consistent with this, our RNA-seq analysis of SG200-infected leaf tissue revealed high expression of Zm*CycD3;1a* and Zm*CycD3;1b,* corroborating previous findings that reported high expression of Zm*CycD3;1a* and Zm*CycD3;1b* in SG200-infected mesophyll HTT cells ^89^.

The pre-RC, consisting of six ORC proteins, CDC6, CDT1, and MCM helicase complex, ensures each daughter cell receives an identical copy of the genome ^53,90^. In *Arabidopsis*, overexpression of At*CDC6a* and At*CDT1a* increased nuclear ploidy in non-dividing cells through extra rounds of endocycle ^54,55^. Similarly, our RNA-seq showed up-regulation of most pre-RC components in SG200-infected leaf tissue, further supporting the role of *hap1* in promoting endoreduplication and linking nutrient availability to cell cycle progression and plant growth.

Cell cycle progression is also controlled by cyclin-dependent kinases (CDKs), which interact with cyclins (CYC) and CDK inhibitors (CKIs) ^91^. In maize, Zm*WEE1*, a negative regulator at the G2/M checkpoint, showed the highest expression in cell cycle arrested maize stamen, indicating its role in endoreduplication ^49^. In tomato, Sl*WEE1* promotes endoreduplication by phosphorylating mitotic CDK/CYCB complex ^46,48^. Knockdown of SI*WEE1* in nematode-induced tomato galls resulted in premature entry into mitosis, smaller gall formation, reduced nematode reproduction, and decreased fruit size ^47^. In *Arabidopsis*, SMR interacts with CDKA/CYCD complexes, preventing mitotic entry and promoting endoreduplication ^50^. In *U. maydis*-infected maize, Zm*SMR1* was shown to be highly expressed in HPT cells and observed increased nuclear size, a parameter linked to cellular endoreduplication, in mesophyll cells infected with SG200 or SG200Δ*see1* strains ^89^. Our RNA-seq analysis revealed up-regulation of Zm*SMR1* and Zm*SMR3* in SG200-infected cells, along with other CDK inhibitors and cyclins involved in endoreduplication. Also, disruption of *hap1* in *U. maydis* resulted in reduced mesophyll nuclear size in maize leaves compared to SG200 infection, reinforcing the role of *hap1* in endoreduplication induction.

ZmRBR1 and ZmRBR3 showed increased phosphorylation in SG200-infected leaf tissue compared to CR-Hap1-infected leaf tissues. RBR proteins, active in their hypophosphorylated state, interact with E2F/dimerization partner (DP) transcription factors during G1, preventing activation of genes required for the G1/S phase transition and inhibiting premature S phase entry ^44,59,60^. Phosphorylation disrupts RBR-E2F interaction, leading to the continuous release of E2F and uncontrolled DNA replication, characteristic of endoreduplicating cells.

### A model of Hap1-mediated modulation of starch metabolism in hypertrophic tumor cells

Young maize seedlings, as photosynthetic sink tissues, require high levels of sugar metabolites for growth. When sugar metabolites such as T6P are low, SnRK1 is activated, initiating the transient starch breakdown by α- and β-amylases (Fig. **7a**). Upon infection, *U. maydis* secretes Hap1 into host cells, elevates T6P levels and creates extensive sink tissues, enhancing nutrient flux that favors fungal colonization. Hap1 interacts with maize SnRK1α, leading to the phosphorylation of SnRK1 targets and up-regulation of enzymes involved in starch biosynthesis. The Hap1-SnRK1 interaction may inhibit T6P-induced immune signaling by promoting the conversion of sugars into starch, thereby disrupting the antagonistic relationship between T6P and SnRK1. This process reprograms transcription required to support starch metabolism, endoreduplication, and suppression of sugar-induced immune responses (Fig. **7b**). On the contrary, in the absence of Hap1, T6P may accumulate in the cell and inactivates SnRK1 to maintain energy homeostasis. Sugar metabolites, which also act as immunosignals, can accumulate excessively in the host cell, potentially triggering immune responses by up-regulating the expression of receptor kinases and transcription factors. This response leads to reduced proliferation of *U. maydis* and a significant reduction in tumor formation (Fig. **7c**).

**Fig. 7).**
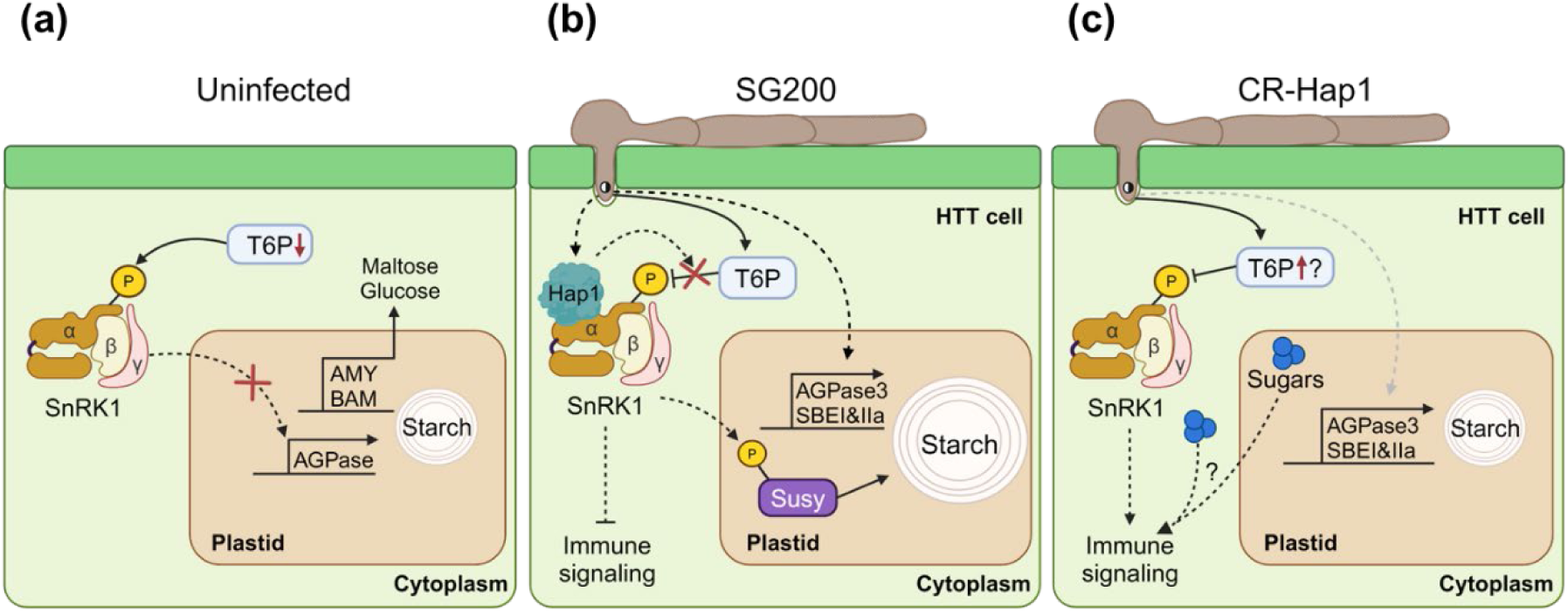
Hap1 disentangles the T6P and SnRK1 antagonism, inducing starch accumulation and hypertrophy in mesophyll tumor cells. **(a)** Activation of SnRK1 under low T6P conditions triggers transient starch breakdown in uninfected maize tissue. **(b)** Effect of Hap1 presence in *U. maydis* on T6P accumulation and SnRK1 activation in infected maize mesophyll tissue. **(c)** Effect of Hap1 absence in *U. maydis* on T6P accumulation and SnRK1 inactivation in maize mesophyll tissue.

Taken together, this study reveals a cooperative role of the Hap effectors, with Hap1 being the major factor in modulating key metabolic processes and cell cycle regulation in maize by interacting with ZmSnRK1. Hap1 binding to ZmSnRK1 rewires carbohydrate signaling and redirect available nutrients to mesophyll cells, ultimately inducing hypertrophy, a hallmarks of *U. maydis* virulence. Together with the previously identified effectors See1 ^7^ and Sts2 ^8^ that induce cell division in bundle sheath cells, discovery of the Hap effectors and their role in mesophyll hypertrophy provides a framework on the molecular basis of *U. maydis*-induced tumorigenesis.

## Material and Methods

### Strains, fungal, and plant growth conditions

Plasmids were cloned in *Escherichia coli* Top10 strains. For transient protein expression in *Nicotiana benthamiana, Agrobacterium tumefaciens* GV3101 was used. *E. coli* and *A. tumefaciens* were grown in dYT-liquid medium (1.6% w/v peptone, 1% w/v yeast extract and 0.5% w/v NaCl) or YT agar plates with appropriate antibiotics at 37°C with shaking at 200 rpm or 28°C, respectively. *U. maydis* was grown in YEPS_Light_ liquid medium (0.4% w/v yeast extract, 0.4% w/v peptone and 2% w/v sucrose) or on PD-agar plates at 28°C with shaking at 200 rpm. *Zea mays* Golden Bantam (GB) or Early Golden Bantam (EGB) plants were grown under controlled glasshouse or phytochamber conditions (16 h light at 28°C and 8 h dark at 22°C). *N. benthamiana* plants were cultivated in a glasshouse (16 h light and 8 h dark at 22°C).

Mutants were generated in *U. maydis* SG200 solopathogenic or effector mutant strains using CRISPR mutagenesis as previously described ^92^. For complementation, the p123 plasmid ^93^, containing an *ip* allele for carboxin (*cbx*) resistance ^94^, was linearized with SspI or AgeI and integrated into the genome via homologous recombination. Single copy integration was confirmed by southern blot (data not shown). Primer details are in Supporting Information Table **S1**.

### Maize infection, and disease scoring

Seven-day-old EGB or GB seedlings were inoculated with *U. maydis* (OD_600_ = 1 for disease scoring and OD_600_ = 3 with 0.1% tween-20 for microscopy, (interactome/total/phospho) proteomics, and RNA-seq). Disease scoring was performed at 12 days post-inoculation (dpi) as previously described ^95^. Disease indexes were assigned as follows: 9 for dead plants, 7 for heavy tumors, 5 for tumors, 3 for small tumors, 1 for chlorosis, and 0 for no symptoms. The disease index was used for assessing statistical significance using Student’s *t*-test with three independent biological replicates. Disease scoring data of maize infection is provided in Supporting Information Data **S1**.

### Split Luciferase complementary assay

*Agrobacterium* carrying the gene of interest was mixed with p19 to OD_600_ = 1 and infiltrated into *N. benthamiana* leaves. 2-3 days post-infiltration, leaves were sprayed with 1 mM D-luciferin (Promega) and incubated in dark for 10 min. Luminescence was detected using a ChemiDoc MP machine (Bio-Rad) from at least three independent plants.

### Co-immunoprecipitation and interaction proteomic analysis

Infected leaves were harvested at 3 dpi and powdered using liquid nitrogen. For protein extraction, 1 ml of powdered leaves was mixed with 1 ml of lysis buffer (50 mM Tris–HCl pH 7.5, 150 mM NaCl, 2 mM EDTA, 10% glycerol, 1% Triton X-100, 5 mM DTT and cOmplete™ protease inhibitor (Roche)). The mixture was incubated on ice for 30 min and centrifuged twice at 4°C with 13 000 *g* for 30 min to obtain protein supernatant. 1ml of the protein supernatant was incubated with anti-Myc or anti-GFP (ChromoTek) magnetic beads at 4°C, 1-2 h with rotation, and washed at least five times. The washed beads were either subjected to mass spectrometry analysis or resuspended in 80 μl of 2× SDS-loading buffer and boiled at 99°C for Western blot analysis. Detection was performed using anti-GFP (Roche), anti-Myc, or anti-HA (Sigma-Aldrich) antibodies. For sample preparation, LC-MS/MS data acquisition, and data analysis of interactome proteomics was performed as previously described for HA enrichment pull-down ^96^. MS/MS spectra were searched by the Andromeda search engine against a combined database containing the sequences from *Z. mays* (Zm-B73-REFERENCE-NAM-5.0) or *U. maydis* (Uniprot) sequences, contaminant proteins, and decoy sequences.

### Total proteome and phosphoproteome analysis

Proteins were extracted with 1ml extraction buffer (8M urea, 20 µl/ml phosphatase Inhibitors (Sigma, P5726-5ML and P0044-5ML, 5 mM DTT) and incubated for 30 min with shaking. Samples were alkylated with 14 mM CAA and quenched with 5 mM DTT. 500 µg of total proteins were diluted to 1 M urea in 100 mM Tris-HCl pH 8.5, 1 mM CaCl_2_, and digested with 5 µg LysC (WAKO) in 50 mM NH_4_HCO_3_ for 4 h at RT. 5 µg trypsin was added to samples and diluted with 100 mM Tris-HCl pH 8.5, 1 mM CaCl_2_. Samples were mixed and incubated overnight at 37°C. Digests were acidified with TFA to 0.5% and desalted using C18 SepPaks (1cc cartridge, 100 mg (WAT023590)). Samples were eluted with 80% ACN/0.1% TFA and dried.

For library samples, 5 µL aliquots of samples were pooled and fractionated using SCX StageTips packed with Empore Cation SPE disks (Empore Cation 2251 material). Samples were fractionated with ammonium acetate gradient (25-500 mM in 20% ACN, 0.5 % FA) into nine fractions, with additional elutions with 1% ammonium hydroxide, 80% ACN and 5% ammonium hydroxide, 80% ACN. Fractions were eluted by centrifugation (5 min, 500 x g), dried, and resuspended in 10 µl A* buffer. For DDA analysis, the remaining 40 µl eluted peptides from the SepPack purification were dried and resuspended in 10 µl A* buffer. Peptide concentration was determined by Nanodrop and samples were diluted to 0.1 µg/µL for measurement.

For phosphopeptide enrichment by metal-oxide chromatography (MOC) (adapted from Nakagami, 2014), peptides were evaporated to 50 µL and diluted with sample buffer (2 ml ACN, 820 µl lactic acid (LA), 2.5 µl TFA / 80% ACN, 0.1% TFA, 300mg/ml LA). MOC tips were loaded with 3 mg TiO_2_ beads (Titansphere TiO_2_, 10 µm (GL Science Inc, Japan, Cat. No. 5020-75010)) in 100 µL MeOH in a C8 micro column. MOC tips were placed onto a 96-well plate (Protein LoBind, (Eppendorf Cat. No. 0030504100) and equilibrated. Samples were loaded onto the tips, centrifuged (10 min at 1000 g), and reloaded for second centrifugation. Tips were washed using solution C and B with centrifugation (1500g, 5 min) and transferred to a fresh 96-well plate containing 100 µl of 20% phosphoric acid for elution of enriched phosphopeptides. Peptide was eluted sequentially with 50 µl elution buffer 1 (5% NH_4_OH) and buffer 2 (10% piperidine), desalted using C18 StageTips ^98^, dried in a vacuum evaporator, and dissolved in 10 µl A* buffer for MS analysis.

### LC-MS/MS data acquisition

Samples were analyzed using an Ultimate 3000 RSLC nano (Thermo Fisher) coupled to an Orbitrap Exploris 480 mass spectrometer with a FAIMS (Field asymmetric ion mobility separation) Pro interface (Thermo Fisher). Peptides were pre-concentrated on an Acclaim PepMap 100 pre-column (Thermo Fisher) using buffer A** (water, 0.1% TFA) at 7 µl/min for 5 min. Peptides were separated on 16 cm frit-less silica emitters (New Objective, 75 µm inner diameter) packed in-house with reverse-phase ReproSil-Pur C18 AQ 1.9 µm resin (Dr. Maisch) and eluted with a segmented linear gradient of 5-95% solvent (80% ACN, 0.1% FA) at 300 nL/min^−1^ over 130 min. Mass spectra were acquired in data-dependent acquisition mode with a TOP_S method using 2 seconds cycle time at resolution of 60,000 FWHM (mass range of 320–1200 m/z) was applied, with a target of normalized AGC 300%. Field asymmetric ion mobility separation (FAIMS) was performed with compensation voltages of −45 and −60 at 1 second cycle time for total proteome and library, and −45 and −65 with 1.2 seconds cycle time (CV-45) and 0.8 seconds cycle time (CV-65) for phosphoproteomics. Precursors were filtered using a 5000-intensity threshold with the MIPS option (MIPS mode = peptide) and selected with a 1.6 m/z isolation window. HCD fragmentation was performed at 30% NCE. MS/MS spectra were acquired with resolution of 15,000 FWHM and target of 75% ion with automated injection time. A fixed first mass of m/z 100 was used for all analyses. Peptides with a charge of +1, >6, or with unassigned were excluded from fragmentation for MS^2^ and dynamic exclusion was applied for 40s to prevent repeated precursor selection.

### Data analysis

Raw data were processed using MaxQuant (v.1.6.3.4, http://www.maxquant.org/) with label-free quantification (LFQ) and iBAQ enabled ^99,100^. MS/MS spectra were searched by the Andromeda search engine against a combined database containing the sequences from *Z. mays* (Zm-B73-REFERENCE-NAM-5.0), contaminant proteins, and decoy sequences. Search parameters included trypsin specificity, up to two missed cleavages, a minimal peptide length of seven amino acids, fixed carbamidomethylation of cysteine, and variable modifications of methionine oxidation and protein N-terminal acetylation. Peptide-spectrum-matches and proteins were retained below FDR of 1%. The match between runs option was enabled.

Statistical analysis of MaxLFQ values was performed using Perseus (v.1.6.14.0). Phosphopeptide intensities were analyzed from the “modificationSpecificPeptides” output. Library and DDA samples were grouped, with the “match from” used for library samples and “match from and to” used for DDA samples. Quantified proteins were filtered for reverse hits and hits “only identified by site” for total proteome and library, and only phospho-modified peptides for phosphoproteomics. MaxLFQ values were log2 transformed and grouped by conditions. Proteins with at least three valid values per condition were processed for mixed imputation by spliting into missing-at-random (MAR) and missing not at random (MNAR) hits based on one valid value per group and splitting the resulting matrices ^101^. The resulting matrix with at least one valid value per group was set as the MAR dataset, and the remaining hits as the MNAR dataset. Missing values were imputed using the “imputeLCMD” R package ^102^ in Perseus: the MAR dataset was imputed using a nearest-neighbor approach (KNN, n=5), and the MNAR dataset using the MinProb option (q=0.01, tune.sigma=1). Two-sample Student’s *t*-tests with a permutation-based FDR of 5% were performed. Results were exported and further processed in Excel.

### RNA preparation and RNA-Seq analysis

For total RNA extraction, leaves were harvested at 3dpi and homogenized in liquid nitrogen using TRIzol Reagent (Invitrogen) followed by DNA removal with the Turbo DNA-Free™ Kit (Ambion Life Technologies™). Three independent biological replicates were subjected to 150-bp paired-end RNA-seq on Illumina NovaSeq 6000 (Illumina) at Novogene (Cambridge, UK). RNA-seq analysis was performed as previously described ^103^.

### Leaf staining, tissue embedding, sectioning and microscopy

For sectioning, leaves were harvested at 6dpi and cut into approximately 2cm (2cm below the infection site). Tissues were embedded as previously described ^7^, molded in Peel-A-Way™, and sectioned transversely into 13 µm slices. For staining, sections were treated with Lugol’s iodide (IKI) solution (Roth) for starch staining and propidium iodide (Sigma-Aldrich) for nuclei staining. Imaging was performed using a Nikon Eclipse Ti inverted microscope with Nikon Instruments NIS-ELEMENTS software (561 nm excitation and at 590–603 nm emission for propidium iodide) or a Thunder microscope with Leica imaging LAS X software.

### GO enrichment and protein-protein interactions (PPI) analysis

GO enrichment analysis was performed using Plaza 5.0 ^104^, with Fisher’s exact test with Bonferroni correction (*P*-value <0.05). Results were grouped into upper hierarchical parent terms using REVIGO ^105^. PPI analysis was performed using the STRING database ^106^.

## Accession numbers

Maize gene sequences were obtained from the maizeGDB database (https://www.maizegdb.org//). The accession numbers of maize SnRK1: SnRK1.1α (Zm00001eb013270) SnRK1.2α (Zm00001eb293240) SnRK1.3α (Zm00001eb094400).

## Acknowledgements

We express our gratitude to Anne Harzen for her valuable assistance in MS sample processing. We acknowledge the funding received from the European Research Council (ERC) under the European Union’s Horizon 2020 research and innovation programme (grant agreement No 771035), as well as the support from the Deutsche Forschungsgemeinschaft (DFG, German Research Foundation) under Germany’s Excellence Strategy-EXC-2048/1-Project ID: 390686111. GD and YJL acknowledge funding by the DFG through project DO1421/3-3. The Max Planck Society is acknowledged for financial support of SCS and HN.

## Author contributions

GD and YJL designed the research. YJL and ME conducted the experiments. GS performed bioinformatics analysis of RNA-Seq data. SCS and HN performed the MS and MS data analysis. YJL and GD wrote the paper with contributions from the other authors.

## Data availability

RNA-seq raw data are publicly available on the NCBI Gene Expression Omnibus with accession number XXXXXX, and the mass spectrometry proteomics data can be accessed in the ProteomeXchange Consortium via the PRIDE repository (Perez-Riverol et al., 2022) under dataset identifier PXD057668 (interaction proteome) and PXD057676 (total and phosphoproteome). Supporting information not provided in the manuscript can be requested from the corresponding author (GD) upon reasonable request.

## Supporting information

### Supplementary Figures

**Fig. S1)** UMAG_00753 is not secreted.

**Fig. S2)** Pull-down/MS analysis of Hap effectors interactomes.

**Fig. S3)** Hap1 is the dominant virulence factor amongst the Hap-effectors.

**Fig. S4)** IP/MS analysis of Hap1 maize interactome and GO enrichment.

**Fig. S5)** GO enrichment analysis of Hap1-CR-Hap2, Hap1-CR-Hap3, and Hap1-CR-Hap2/3 in *Zea mays*.

**Fig. S6)** Differentially expressed genes in comparison of *Ustilago maydis*-infected (SG200 or CR-*hap1*) to mock-infected treatments.

**Fig. S7)** The absence of Hap1 affects the expression of defense response related genes.

**Fig. S8)** Venn diagram analysis of unique and overlapping proteins between phosphoproteomics and total proteomics in SG200 vs. mock and CR-Hap1 vs. mock.

### Supplementary Data

**Data S1)** Scoring of disease symptoms in infected maize seedlings at 12 days post-inoculation.

**Data S2)** Secreted effector proteins identified as potential Hap1 interactors in Hap1 IP-MS data from *U. maydis*.

**Data S3)** Secreted effector proteins identified as potential Hap2 interactors in Hap2 IP-MS data from *U. maydis*.

**Data S4)** Secreted effector proteins identified as potential Hap3 interactors in Hap3 IP-MS data from *U. maydis*.

**Data S5)** List of kinases, phosphatases, trehalose-6-phosphate synthases, and starch biosynthetic enzymes identified in the mass spectrometry data for Hap1-CR-Hap2, Hap1-CR-Hap3, and Hap1-CR-Hap2/3 in Zea mays.

**Data S6)** Differentially expressed genes of SG200-infected vs. mock-infected maize.

**Data S7)** Differentially expressed genes of CR-Hap1-infected vs. mock-infected maize.

**Data S8)** Differentially expressed genes of CR-Hap1-infected vs. SG200-infected maize.

**Data S9)** List of upregulated defense-related differentially expressed genes.

**Data S10)** List of upregulated differentially expressed genes involved in cell cycle regulation.

**Data S11)** List of upregulated differentially expressed genes involved in starch biosynthesis.

**Data S12)** Gene ontology analysis of increased phosphoproteins in SG200-infected vs. mock-infected maize.

**Data S13)** Gene ontology analysis of increased phosphoproteins in CR-Hap1-infected vs. mock-infected maize.

**Data S14)** Overview of the SnRK1-dependent consensus motif at P-5 and P+4 relative to phosphorylated Ser/Thr residues.

### Supplementary Tables

**Table S1)** Oligonucleotides and plant infection disease symptom scoring in this study.

**Table S2)** Mass spectrometry data of Hap1 in *U. maydis*.

**Table S3)** Mass spectrometry data of Hap2 in *U. maydis*.

**Table S4)** Mass spectrometry data of Hap3 in *U. maydis*.

**Table S5)** Mass spectrometry data of Hap1 in *Z. mays*.

**Table S6)** Mass spectrometry data of Hap1-CR-Hap2 in *Z. mays*.

**Table S7)** Mass spectrometry data of Hap1-CR-Hap3 in *Z. mays*.

**Table S8)** Mass spectrometry data of Hap1-CR-Hap2/3 in *Z. mays*.

**Table S9)** Shotgun (total proteomic) data of Mock, SG200, and CR-Hap1 in *Z. mays*.

**Table S10)** Phosphoproteomic data of Mock, SG200, and CR-Hap1 in *Z. mays*.

