## Supplemental Figures for "*Ustilago maydis* disrupts carbohydrate signaling networks to induce hypertrophy in host cells"

**Supporting information: Tables S1 – S8 and Datasets S1 – S12**

**Supporting Figures**

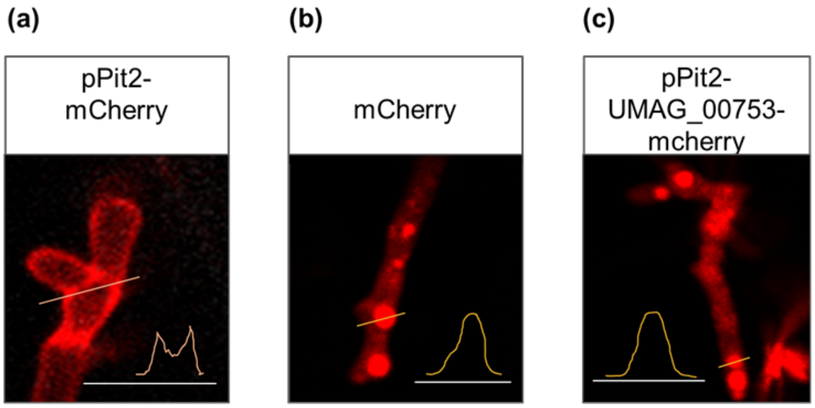


**Fig. S1) UMAG_00753 is not secreted.** Confocal laser scanning microscopy images depicting secretion of UMAG_00753 upon maize infection at 2 days post infection. Fluorescent signals were observed for the following *U .maydis* strains: (**a**) SG200-CR-*pit2*_p*Pit2*::*Pit2-mCherry*, (**b**) SG200-p*Pit2*::*mCherry*, or (**c**) SG200-CR-*pit2*_p*Pit2*::*UMAG_00753-mCherry*. SG200-CR-*pit2*_p*Pit2*::*Pit2-mCherry* show fluorescence signal at the hyphal and tips, while SG200-p*Pit2*::*mCherry* and SG200 *pit2*_p*Pit2*::*UMAG_00753-mCherry* show signals within the hyphae. Fluorescence intensity profiles of detected signals along the orange lines are illustrated at the bottom of each respective images. Representative images from two independent experiments are displayed. Scale bar =50µm

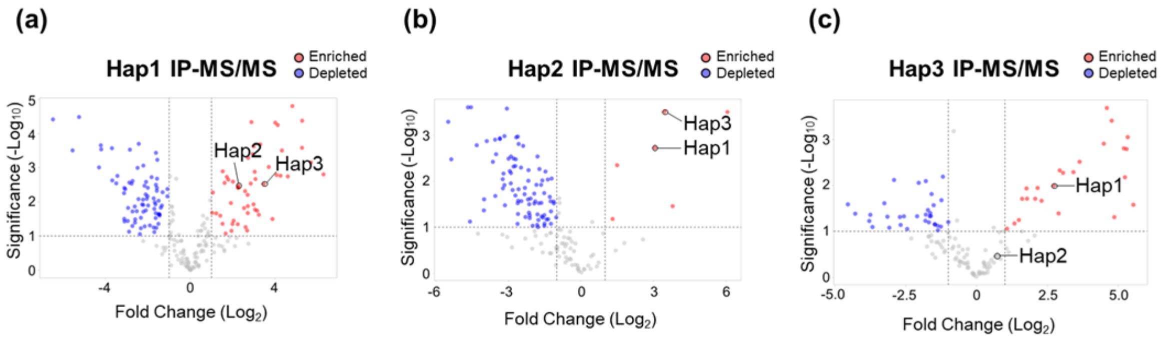


**Fig. S2) Pull-down/MS analysis of Hap effectors interactomes.** Volcano plots of displaying Hap effector interacting proteins detected by LC/MS. The fold change (FC) values were calculated by dividing the LFQ intensities of protein peptides in (**a**) Hap1, (**b**) Hap2, or (**c**) Hap3 compared to the control. The x-axis represents $Log_2FC$, and the y-axis represents high statistical significance ($-Log_{10}$ of *P-values*). Proteins with a $Log_2FC$ of >1.0 or <-1 and *P*< 0.05, as determined by Student's t-test are shown. Red and blue dots indicate up- and down-regulated proteins, respectively, while gray dots indicate proteins with no significant changes.

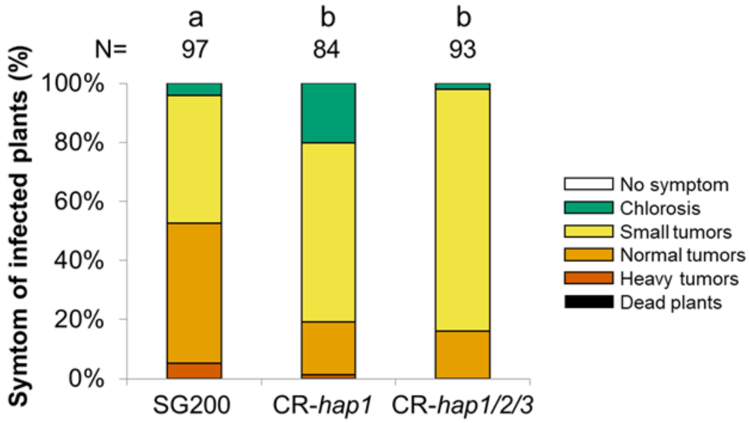


**Fig. S3**) **Hap1 is the dominant virulence factor amongst the Hap-effectors.** The disease index is presented as the mean from three biological replicates. Statistical significance was determined using Tukey's HSD post hoc test following a one-way ANOVA, $P<0.05$. Different letters indicate significant differences between treatments.

(a)

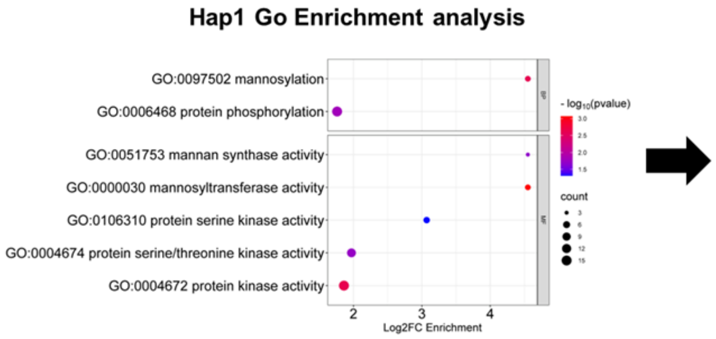


(b)

| Description |
|---|
| RING-type E3 ubiquitin transferase |
| RING-type E3 ubiquitin transferase |
| Serine/threonine-protein kinase SAPK5 |
| OSJNBa0016N04.16-like protein |
| Cysteine-rich receptor-like protein kinase 6 |
| Outer arm dynein light chain 1 protein |
| ZmSnRK3.12 |
| Protein kinase superfamily protein |
| Mitogen-activated protein kinase kinase 3 |
| Putative MAP kinase family protein |
| Calcium-dependent protein kinase 7 |
| ZmSnRK1α1 |
| ZmSnRK1α2 |
| ZmSnRK1α3 |
| ZmSnRK1α4 |

**Fig. S4) IP/MS analysis of Hap1 maize interactome and GO enrichment.** (**a**) Gene ontology (GO) enrichment analysis was performed on Top 150 maize proteins interacting with Hap1 using PLAZA 5.0. GO terms were grouped into higher hierarchical categories and summarized with REVIGO. (**b**) A list of proteins showing all biological processes identified in Fig. S4a.

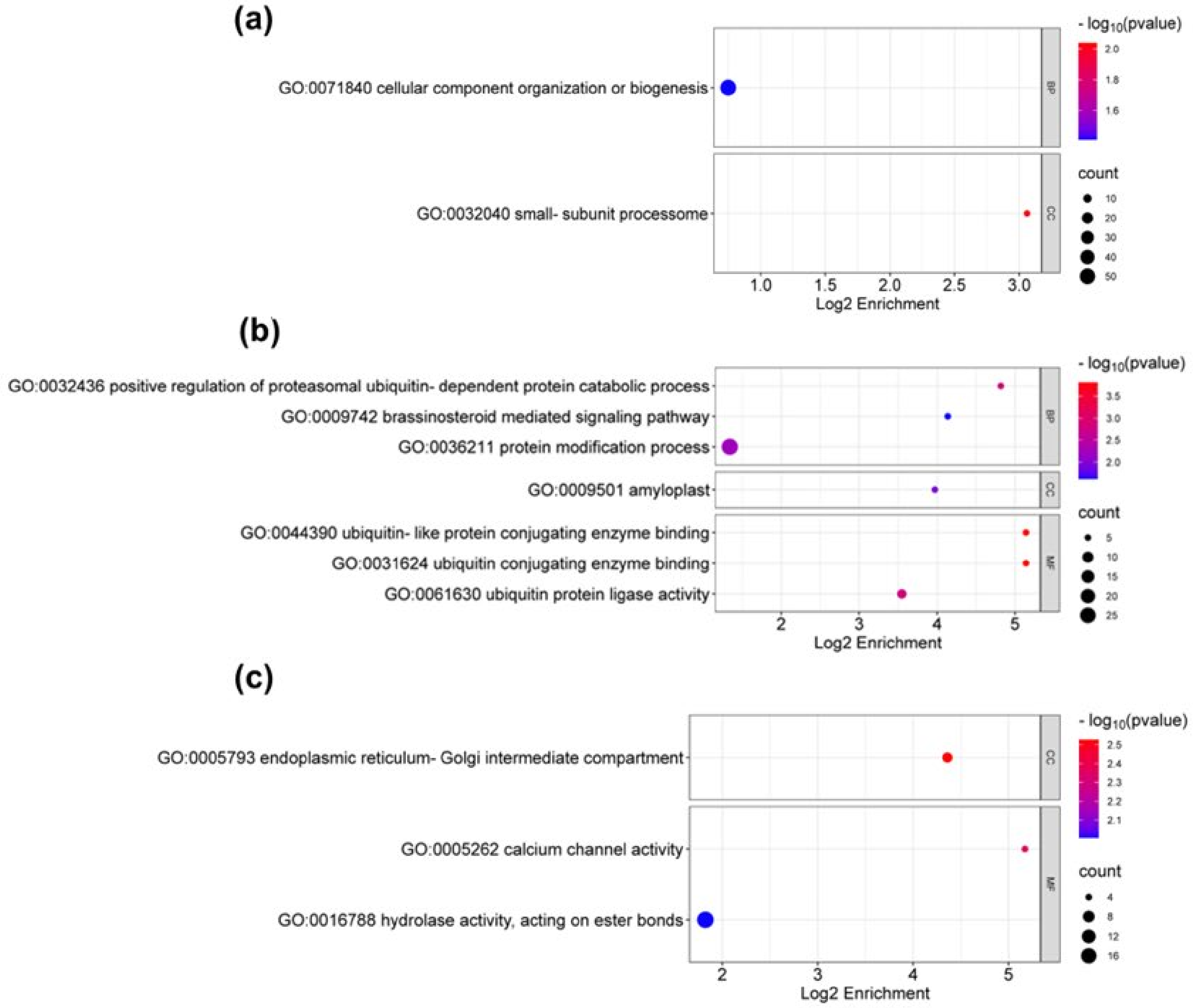


**Fig. S5) GO enrichment analysis of Hap1-CR-Hap2, Hap1-CR-Hap3, and Hap1-CR-Hap2/3 in *Zea mays.* (a**) GO enrichment analysis of proteins enriched in Hap1-CR-Hap2. (**b**) GO enrichment analysis of proteins enriched in Hap1-CR-Hap3. (**c**) GO enrichment analysis of proteins enriched in Hap1-CR-Hap2/3. $Log_2FC>1$ or <-1 and $P<0.05$.

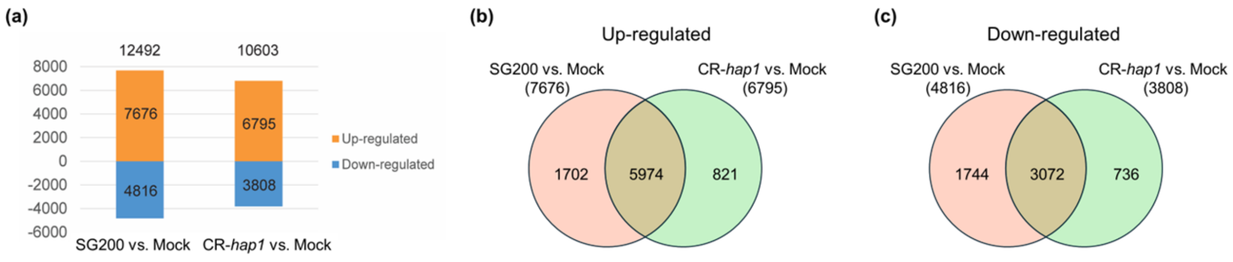


**Fig. S6) Differentially expressed genes in comparison of *Ustilago maydis*-infected (SG200 or CR-*hap1*) to mock-infected treatments.** (**a**) Pairwise bar graph depicting the number of up- and down-regulated DEGs in *U. mayids*-infected (SG200 or CR-*hap1*) vs. mock-infected treatments. DEGs = $Log_2FC>1$ or <-1 and FDR <0.05. (**b-c**) Venn diagram depicting the number of commonly shared and unique DEGs of up- and down-regulated genes in *U. mayids*-infected (SG200 or CR-*hap1*) vs. mock-infected treatments, respectively.

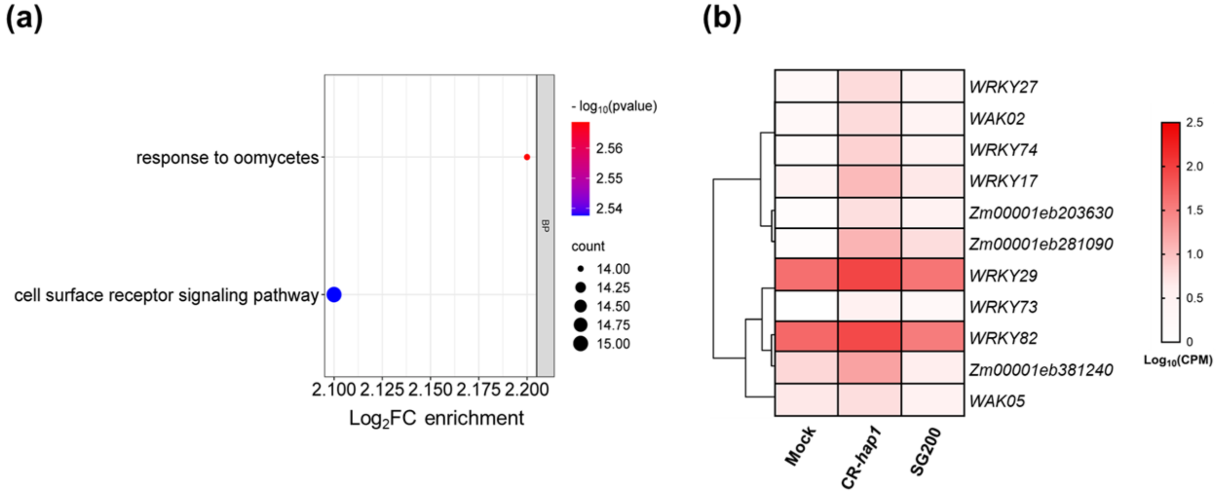


**Fig. S7) The absence of Hap1 affects the expression of defense response related genes** (**a**) GO enrichment analysis of DEGs up-regulated in CR-*hap1.* **(b)** Heatmap of DEGs related to plant defense response, specifically related to WRKY transcription factors, WAKs. DEGs = $Log_2FC>1$ or <-1 and FDR <0.05.

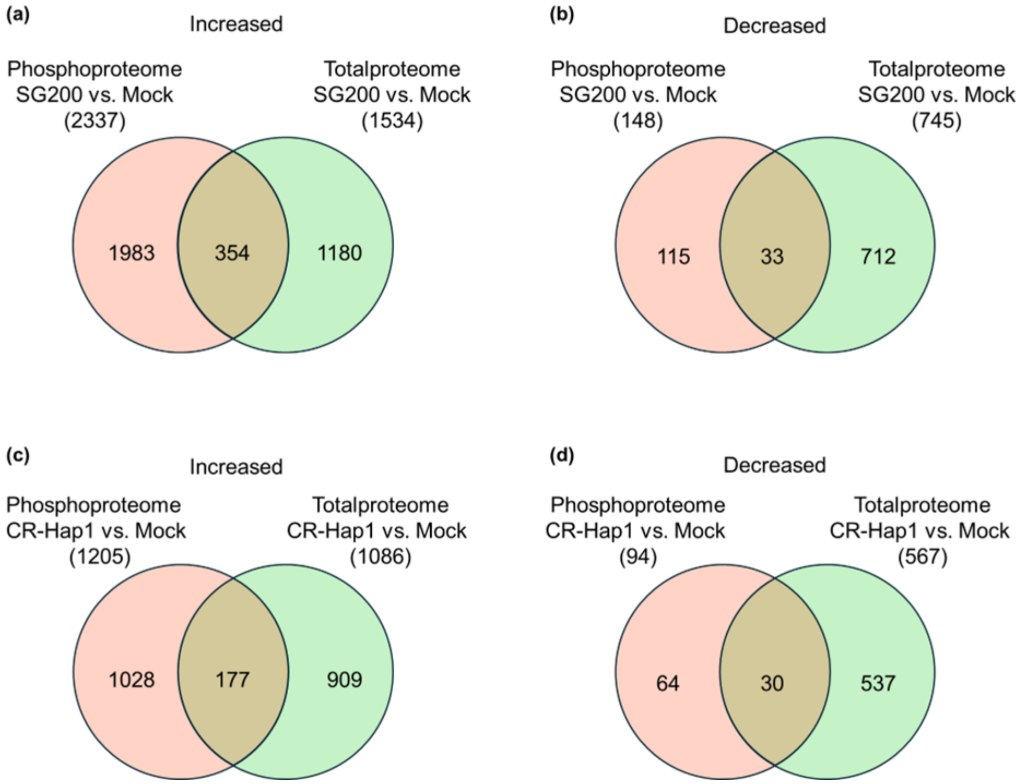


**Fig. S8) Venn diagram analysis of unique and overlapping proteins between phosphoproteomics and total proteomics in SG200 vs. mock and CR-Hap1 vs.mock.** The Venn diagram depicts (**a**) increased phosphoproteins and enriched total proteins in SG200 vs. mock. (**b**) decreased phosphoprotein and depleted total proteins in SG200 vs. mock. (**c**) increased phosphoproteins and enriched total proteins in CR-Hap1 vs. mock (**d**) decreased phosphoprotein and depleted total proteins in CR-Hap1 vs. mock
